# Nonergodicity and Simpson’s paradox in neurocognitive dynamics of cognitive control

**DOI:** 10.1101/2024.07.05.602273

**Authors:** Percy K. Mistry, Nicholas K. Branigan, Zhiyao Gao, Weidong Cai, Vinod Menon

## Abstract

Nonergodicity and Simpson’s paradox present significant, yet underappreciated challenges in neuroscience. Leveraging brain imaging and behavioral data from over 4,000 children and a Bayesian computational model of cognitive dynamics, we investigated brain-behavior relationships underlying cognitive control at both between-subjects and within-subjects levels. Strikingly, we observed a reversal of associations of inhibitory control brain activations with dynamic behavioral measures when comparing between-subjects and within-subjects analyses, revealing the nonergodic nature of these processes. This nonergodicity was pervasive throughout the brain but most pronounced in the salience network. Additionally, within-subjects analysis uncovered dissociated brain representations of reactive and proactive control processes, as well as distinct brain-behavior associations for individuals who adaptively versus maladaptively regulated cognitive control. Our findings offer insights into dynamic neural mechanisms of cognitive control during a critical developmental period. This work highlights the importance of embracing nonergodicity in human neuroscience, with implications for both theoretical understanding and applications to AI and psychopathology.

## Introduction

Since the nineteenth century, the concept of ergodicity has been central to physics^1–3^. However, only recently have other disciplines such as economics^4^ and psychology^5,6^ begun to examine how their own theories and findings rest upon assumptions of ergodicity. Nonergodicity refers to a system’s behavior where time averages and ensemble averages do not coincide^1^. In psychology, a classic case of nonergodicity is Simpson’s paradox observed in the context of the speed-accuracy tradeoff: speed and accuracy are often positively correlated between individuals (faster people are more accurate)^7^, but are negatively correlated within individuals (when an individual tries to respond faster, their accuracy decreases)^8^. This striking divergence between group-level patterns and individual-level dynamics underscores the nonergodic nature of behavioral processes.

The implications of nonergodicity are profound. Analyzing data from a single individual over time can yield different conclusions than analyzing data across multiple individuals at a single point in time^5,6,9^. This challenges traditional assumptions about the generalizability of findings from between-subjects analyses to individual processes^5,9–13^ changing how we interpret and generalize results in the life sciences. While progress has been made in understanding nonergodicity within behavioral contexts^5,6,14–17^, its application to human brain function remains unexplored. This gap in research is particularly striking given that most human neuroscience studies of brain-behavior relations rely heavily on between-subjects analysis. If neurocognitive processes are indeed nonergodic, considering within-subjects analysis of neuroscientific data could profoundly enhance our understanding of brain function.

Nonergodicity in neurocognitive function may manifest as a divergence between group-level and individual-level brain-behavior relationships. Specifically, the association between neural activity and behavior may differ substantially when examined between subjects after averaging over time (**Figure 1a**) compared to within subjects over time (**Figure 1b**), potentially leading to a neurocognitive manifestation of Simpson’s paradox (**Figure 1c**)^11,16^. This phenomenon has profound implications for our understanding of brain function and cognition. Group-level relationships may obscure the dynamic processes occurring within each individual, leading to incomplete or even misleading conclusions about the neural mechanisms underlying cognitive processes^5,6,14–17^.

**Figure 1.**
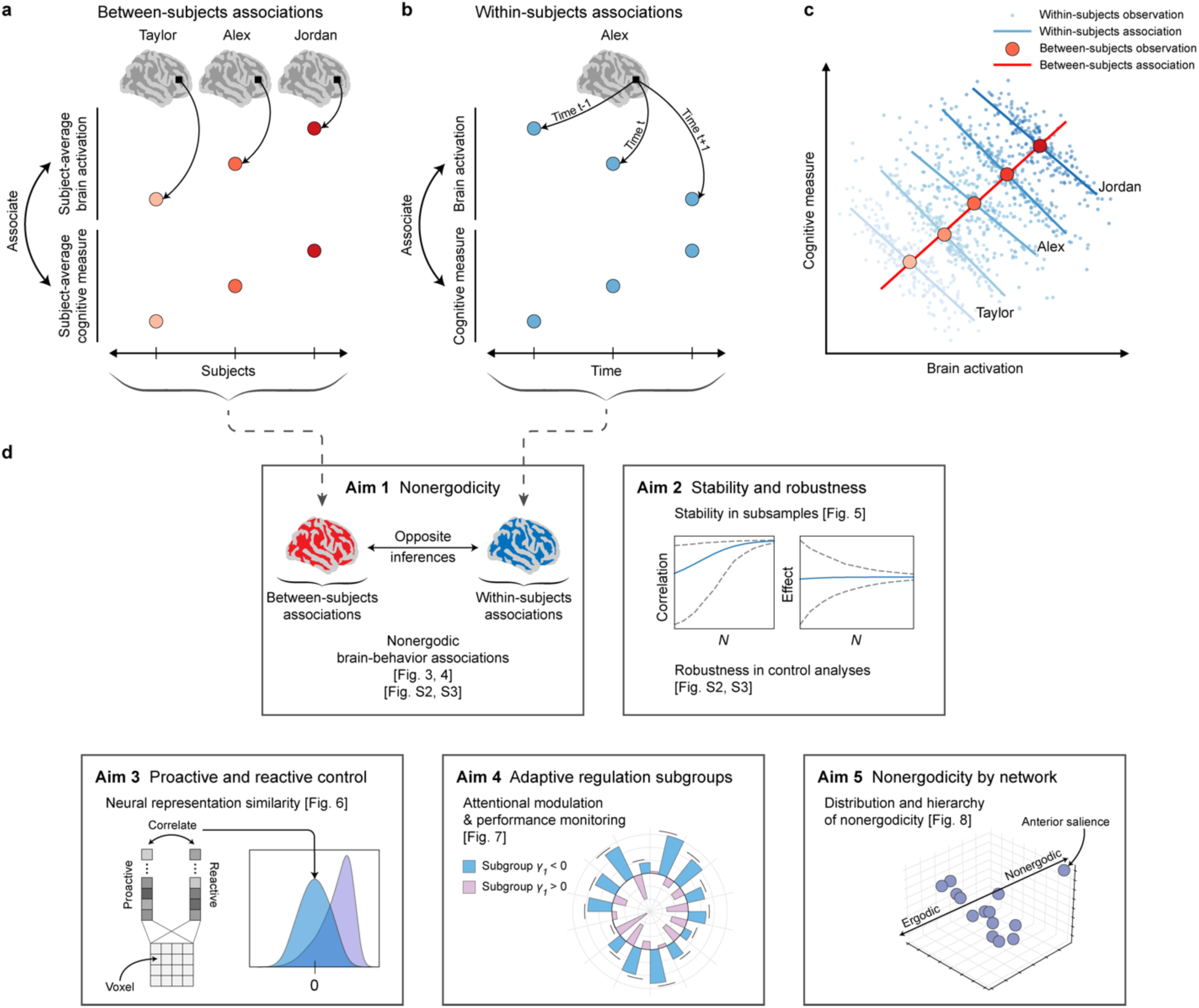
Conceptual overview of the study and key findings. The figure illustrates the methodology for between-subjects and within-subjects analyses, the concept of nonergodicity, and previews the main results. **a,** Between-subjects analysis. Subject-average brain activation in each voxel is correlated with a subject-average cognitive measure across the population. **b,** Within-subjects analysis. For each individual, the time-series of brain activity in each voxel is associated with the time-series of a cognitive measure. **c,** Simpson’s paradox. Simpson’s paradox occurs when associations disagree between subjects and within subjects; it exemplifies nonergodicity in the behavioral sciences. **d,** Study aims. We examined brain-behavior associations for nonergodicity; tested our within-subjects results for stability and robustness; used these results to probe the brain implementations of proactive and reactive control and adaptive regulation of inhibitory control; and investigated how nonergodicity varied by brain network. Nonergodic patterns in brain-behavior associations were consistently observed, revealing that group-level (between-subjects) and individual-level (within-subjects) associations yield divergent results for inhibitory control processes. This challenges the common assumption that findings from such group-level analyses can be directly applied to understand individual- level cognitive processes.

The integration of nonergodic principles into human neuroscience not only promises to advance theoretical understanding but also holds practical implications for tailoring therapeutic strategies to better address individual neurological and cognitive differences. This approach could lead to a more precise and accurate understanding of brain function, paving the way for personalized interventions in clinical neuroscience and psychiatry. There is a pressing need to integrate nonergodic principles into the study of human cognition and brain function, particularly in complex cognitive domains such as inhibitory control, where individual variability and dynamic processes play crucial roles.

To address critical gaps in our knowledge of nonergodicity in neurocognitive processes, we leveraged a large sample (N∼4,000) from the Adolescent Brain Cognitive Development (ABCD) study, a multisite longitudinal study of brain development and behavior in children and adolescents^18^. We used behavioral and brain imaging data to examine the neural mechanisms of cognitive control at both the between-subjects and within-subjects levels, aiming to elucidate the nonergodic nature of these processes and its implications for our understanding of brain function and cognition.

An important component of human cognition is inhibitory control, which is the ability to withhold or cancel maladaptive actions, thoughts, and emotions, and is a fundamental component of goal-directed behavior^19,20^. This critical cognitive function allows individuals to navigate complex environments, adapt to changing circumstances, and maintain focus on long-term goals in the face of immediate temptations or distractions^21^. Given its central role in cognitive control, understanding the neural mechanisms underlying inhibitory control has been a primary focus of cognitive neuroscience research^22^. Inhibitory control engages a distributed network of cortical and subcortical regions. Previous studies have implicated the involvement of the salience network, particularly the anterior insula and dorsal anterior cingulate cortex, in detecting and processing relevant stimuli and coordinating neural resources for inhibitory control^23–27^. Additionally, the inferior frontal gyrus, presupplementary motor area, and basal ganglia have been shown to play crucial roles in implementing response inhibition^28–34^. While the dynamic interplay between these cortical and subcortical regions is known to underpin successful inhibitory control, our understanding of how these neural processes relate to moment-to-moment fluctuations in cognitive control at the individual level remains limited.

The stop signal task (SST) is a widely used paradigm to assess inhibitory control. In this task, participants are instructed to respond quickly to a primary “go” stimulus but to withhold their response when an infrequent “stop” signal appears^20^. The ability to inhibit responses is quantified using the stop signal reaction time (SSRT), which estimates the latency of the stopping process^20,21^. Conventionally, studies investigating the neural basis of inhibitory control have relied on between-subjects analyses of SSRT, comparing brain activity across individuals with varying abilities to inhibit responses. These analyses typically correlate individual differences in SSRT with differences in task-related brain activation patterns^24,35–41^. Such approaches aggregate data at the individual level and make inferences about the neural mechanisms of inhibitory control based on group-level relationships. However, this conventional approach implicitly assumes ergodicity—specifically that the average trend in the group reflects the trend within each individual^6,16,42^. If this assumption does not hold, then inferences made from group-level associations cannot be generalized to explain the cognitive processes occurring within each individual^11,16^.

However, testing the ergodicity assumption in inhibitory control studies has been challenging, particularly when using paradigms like the SST. This is because traditional models do not provide trial-wise SSRT estimates, instead requiring a substantial number of trials for a single estimate^43^. This limitation has constrained researchers’ ability to examine within-subjects dynamics directly, making it difficult to verify whether group-level relationships truly reflect individual-level processes.

To address these limitations and capture the dynamic nature of inhibitory control, we developed the Proactive Reactive and Attentional Dynamics (PRAD) model of SST behavior. This computational model extends beyond conventional approaches to the SST by incorporating proactive and reactive control mechanisms^44^. The PRAD model allows us to infer dynamic, trial-level measures corresponding to elemental neurocognitive subprocesses for each subject, including SSRTs and measures of proactive delaying based on stop signal anticipation. PRAD parameters characterize the latent cognitive constructs governing action execution and inhibition in the SST, providing a more comprehensive view of inhibitory control processes than traditional models.

The PRAD model dissociates observed behavior into elemental latent neurocognitive subprocesses, that can characterize multiple dynamic measures of within-subjects variation. This is critical to conducting within-subjects analysis and evaluating ergodicity, since behavioral variability within a cognitive task is often governed by the complex interplay of multiple neurocognitive subprocesses, each of which may potentially have distinct underlying brain dynamics. By considering the various sequential dependencies and compensatory, competing, and interacting neurocognitive mechanisms underlying cognitive control, the PRAD model dissociates elemental processes and improves the interpretability of inferred within-subjects brain-behavior relationships. Specifically, the PRAD model can infer non-random and sequentially dependent trial-level variability in the probability and duration of proactive delayed responding^45^, as well as in the effect of attentional modulation on the reactive SSRT.

We combined task fMRI data with the PRAD model to investigate the relationship between task-evoked brain responses and dynamic inhibitory control processes at both between-subjects and within-subjects levels. This approach allows us to directly compare inferences made from traditional group-level analyses (group-level associations) with those derived from a fine-grained examination of within-individual variation in neurocognitive dynamics. We leveraged the dynamic representations of subjects’ behaviors from the PRAD cognitive model alongside simultaneous task fMRI data to probe how moment-to-moment variations in brain activity relate to fluctuations in inhibitory control processes. This analysis strategy promotes a richer understanding of the neural basis of inhibitory control, potentially revealing insights that would be obscured in conventional group-level analyses.

We had five interconnected aims (**Figure 1d**). First, we examined how key parameters of reactive and proactive control^46^ from the PRAD model related to brain activity at both between-subjects and within-subjects levels, identifying networks showing significant associations. By comparing these levels, we show the limitations of assuming ergodicity in the study of inhibitory control and underscore the importance of considering within-subjects variability. Second, motivated by the replicability crisis in neuroscience research, we assessed the stability and robustness of our findings. We examined the stability of our within-subjects findings as a function of the sample and sample size through bootstrap resampling. Then, we tested whether our finding of nonergodicity was robust to alternative between-subjects analyses using different measures of brain activation and robust to the use of a directly observed behavioral measure in place of latent cognitive model parameters. Third, we probed the brain’s implementation of proactive and reactive control processes underlying inhibitory control. Proactive control involves the anticipation and preparation for stopping, while reactive control involves the actual implementation of response inhibition^46^. By comparing the brain representations of these processes, we improve understanding of how their neural underpinnings relate. Fourth, we identified subgroups of within-subjects results associated with individual differences in adaptive regulation of inhibitory control. Finally, to characterize the spatial distribution of nonergodicity across the brain, we quantified and compared nonergodicity levels across different functional networks. We defined a network’s degree of nonergodicity as the proportion of individuals whose within-subjects brain-behavior associations exhibited a direction opposite to that observed in the between-subjects analysis. This approach allowed us to identify brain networks with significant levels of nonergodicity.

Our findings demonstrate a stark contrast between traditional between-subjects analyses and novel within-subjects analyses that account for dynamic cognitive processes. Results not only highlight the limitations of assuming ergodicity in cognitive control research but also provide a more precise understanding of the neural mechanisms underlying inhibitory control at the individual subject level. Our within-subjects analysis of neurocognitive dynamics reveals insights that are either obscured by or entirely unavailable from conventional between-subjects approaches. These discoveries have profound implications for both basic neurocognitive research and clinical applications.

## Results

### Dynamic cognitive process model of behavior

We used data from the SST (**Figure 2a**) to investigate dynamic cognitive processes underlying cognitive control in a large sample (*N*∼4000) from the baseline visit of the ABCD study. We developed PRAD, a dynamic cognitive process model^44^ that provides a multidimensional perspective of the elemental cognitive processes governing the reactive and proactive dynamics involved in inhibitory control. This model incorporates latent dynamics that respond to internal cognitive states (endogenous variables) and external environmental contingencies (exogenous variables), with interaction between these dynamics governed by latent trait measures. The latent trait (individual-level) and dynamic (trial-level) measures are simultaneously inferred within a hierarchical Bayesian framework (**Figure 2b**). For each subject and trial, the model infers a measure of reactive inhibitory control, SSRT, and 2 measures of proactive control, probability of proactivity and proactive delaying (**Figure 2c-d**). SSRT measures how long it takes for a subject to inhibit a response, probability of proactivity measures whether they use a strategy of proactive delaying, and proactive delaying measures the length of proactive delays when a proactive strategy is utilized^45,47^. Importantly, all 3 of these parameters are inferred for each trial (**Figure 2e**). **Supplementary Figure S1** provides a detailed illustration of the model. Validation of the PRAD model can be found in a separate study^44^.

**Figure 2.**
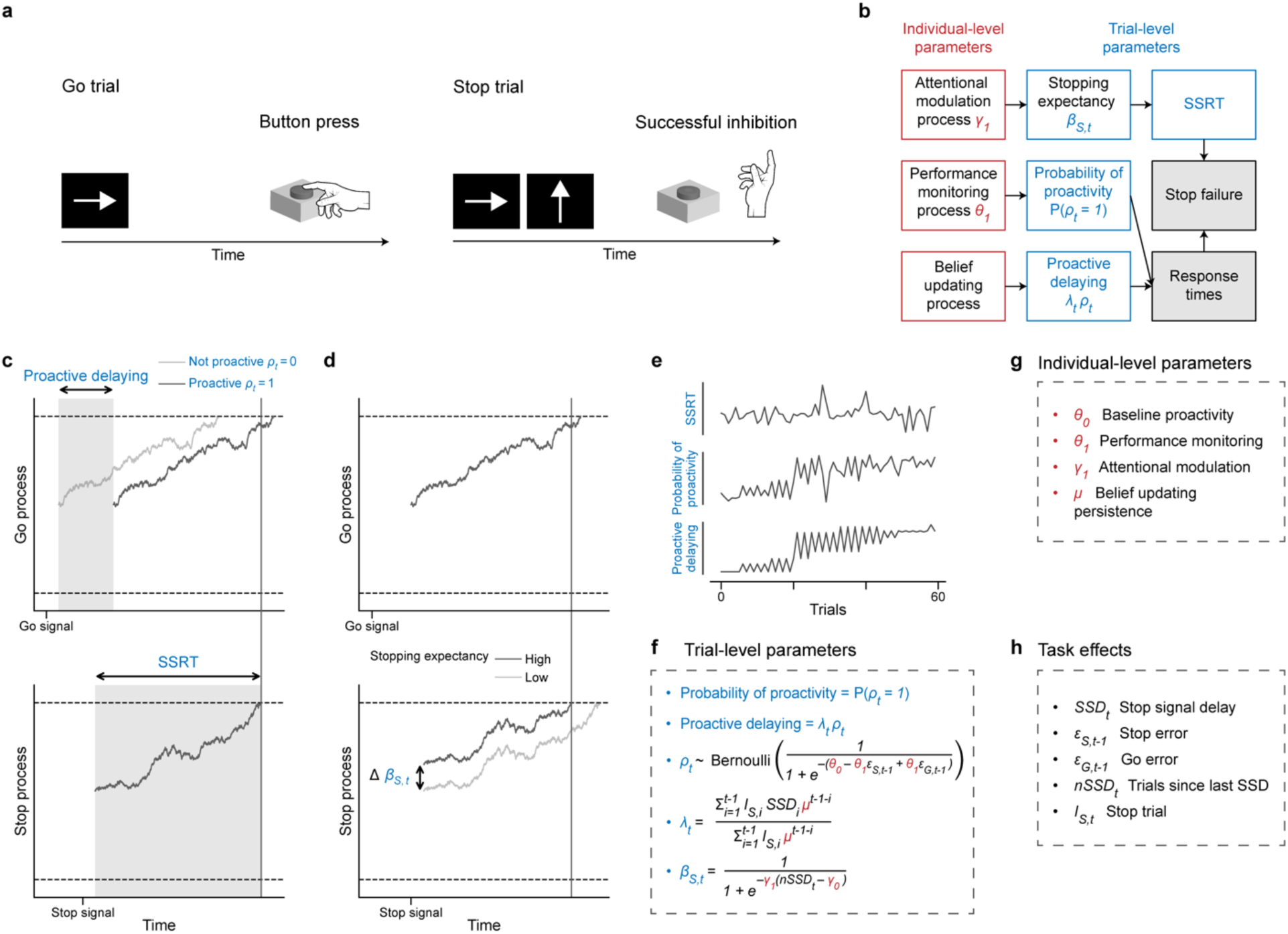
Computational modeling of inhibitory control dynamics in the stop signal task. **a**, Stop signal task. On go trials subjects should respond by pressing a button to indicate the arrow direction, and on stop trials should inhibit their response when the stop signal appears. **b,** Computational model schematic. The computational model infers individual-level and trial-level parameters that measure inhibitory control from observed stop failure and response times. SSRT is the time it takes for the stop process to complete. Probability of proactivity is the probability that a subject uses a proactive control strategy. Proactive delaying is the delay in initiating the go process when a subject uses a proactive control strategy. **c,d,** Within-trial dynamics. **c,** Increased proactivity improves stop trial performance through its effect on the go process. **d,** Increased stopping expectancy improves stop trial performance through its effect on SSRT. **e,** Trial-by-trial dynamics. Illustration of how SSRT, probability of proactivity, and proactive delaying might vary over time within an individual. **f-h,** Key model quantities and their relations. The computational model allows for a detailed, dynamic analysis of inhibitory control processes with trial-level temporal resolution, enabling the investigation of within-subjects variability and nonergodic patterns in cognitive control.

### Between-subjects and within-subjects analysis of brain-behavior associations

We examined how 3 key parameters of the PRAD model that were inferred at a trial level (dynamic measures)—SSRT, probability of proactivity, and proactive delaying—related to brain activity at both between-subjects and within-subjects levels (**Figure 3**). To investigate brain-behavior relationships in inhibitory control, we conducted both between-subjects and within-subjects analyses using fMRI data from the stop signal task (SSRT and probability of proactivity *N* = 4469; proactive delaying *N* = 4176). For the between-subjects analyses, we Pearson correlated subject-average brain activation during successful stopping (correct stop versus correct go activation) with subject-average SSRT, probability of proactivity, and proactive delaying across participants^24,37–41^. For the within-subjects analyses, we regressed the fMRI signal on the parameters using fMRI general linear models with SSRT, probability of proactivity, and proactive delaying included as regressors. We thus modeled the fMRI signal as a combination of static trial-type effects and dynamic effects proportional to trial-by-trial variations in the cognitive model parameters. This allowed us to examine how fluctuations in cognitive processes covaried with brain activity within each individual, while adjusting for stimulus types. These complementary approaches enabled us to compare group-level and individual-level trends in brain-behavior relationships.

**Figure 3.**
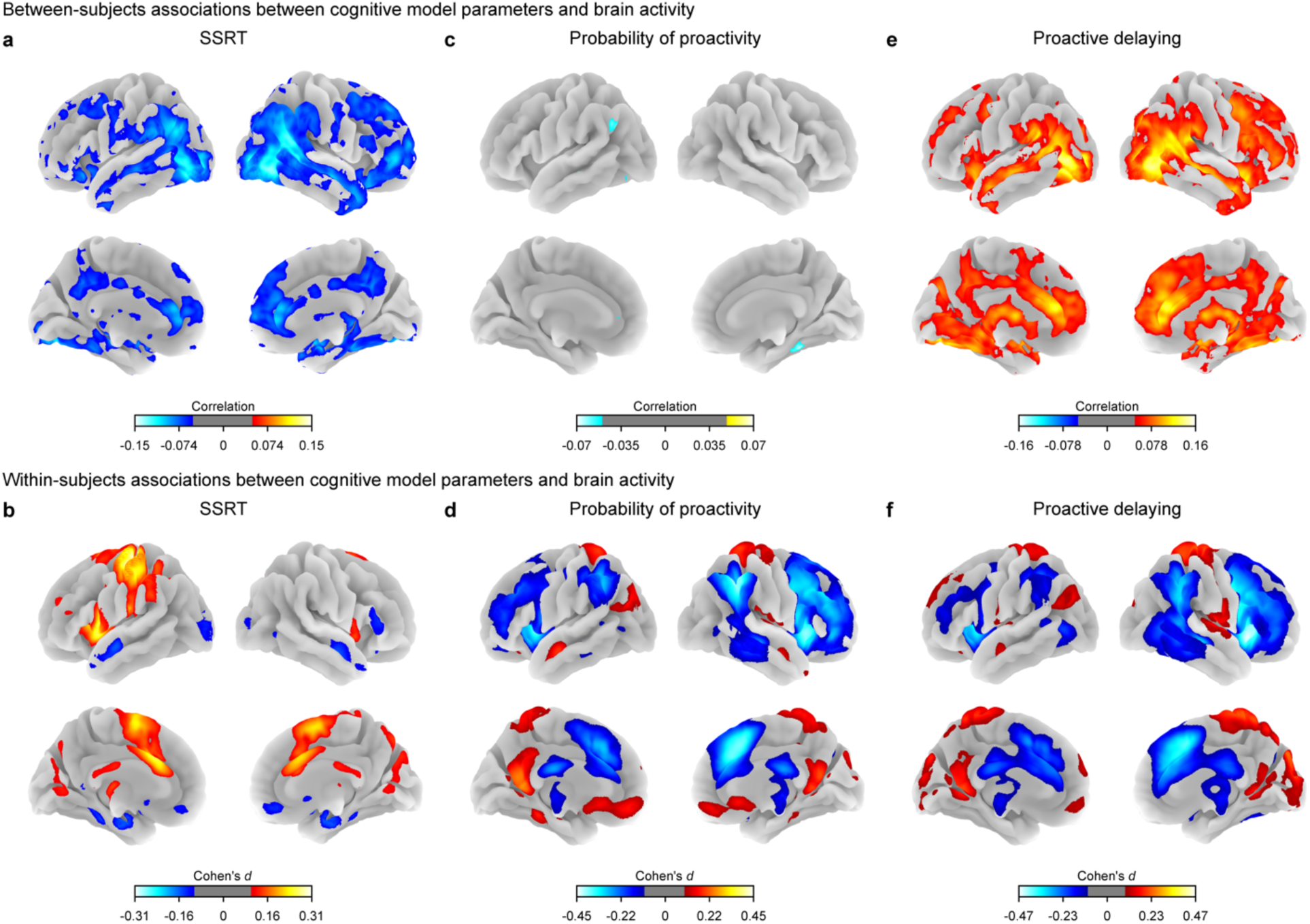
Divergence of between-subjects and within-subjects brain-behavior associations in inhibitory control. **a,c,e,** Between-subjects analysis. Whole-brain correlation maps showing associations between subject-average brain activation (correct stop versus correct go activation) and subject-average cognitive model parameters: SSRT **(a)**, probability of proactivity **(c)**, and proactive delaying **(e)**. Thresholded at Pearson *r* ≥ 0.05. **b,d,f,** Within-subjects analysis. Whole-brain Cohen’s *d* maps showing associations between trial-by-trial brain activity and cognitive model parameters: SSRT **(b)**, probability of proactivity **(d)**, and proactive delaying **(f)**. SSRT associations were computed on stop trials; probability of proactivity and proactive delaying associations were computed on all trials. Thresholded at Cohen’s *d* ≥ 0.1. For both between- and within-subjects analyses: SSRT and probability of proactivity *N* = 4469; proactive delaying *N* = 4176. Striking differences were observed comparing between-subjects and within-subjects associations across multiple brain regions. This divergence provides evidence for nonergodicity in inhibitory control processes, challenging the assumption that group-level findings can be directly applied to understand individual-level cognitive dynamics.

#### SSRT

Between subjects, SSRT showed widespread negative correlations with brain activity, including in frontal, parietal, and temporal areas, and in regions implicated in cognitive control (**Figure 3a**). This suggests that individuals with faster inhibitory responses (lower SSRT) show greater activation in these regions during successful stopping, replicating previous findings^41^. In contrast, within subjects, SSRT showed positive associations with brain activity, particularly in frontal and parietal regions (**Figure 3b**). This suggests that on trials where individuals have slower inhibitory responses, they show increased activation in these areas.

#### Probability of proactivity

Between subjects, probability of proactivity showed minimal correlations with brain activity (**Figure 3c**). In contrast, within subjects, probability of proactivity showed negative associations in parietal, temporal, and lateral frontal regions, and positive associations in default mode network regions (**Figure 3d**).

#### Proactive delaying

Between subjects, proactive delaying showed widespread positive correlations with brain activity, particularly in frontal and parietal regions (**Figure 3e**). This indicates that individuals who engage in more proactive delaying exhibit higher activation in these areas during successful stopping. In contrast, within-subjects, proactive delaying showed negative associations in frontal and parietal cortex (**Figure 3f**).

These findings reveal a striking divergence of between-subjects and within-subjects brain-behavior relationships in inhibitory control. The reversal of association directions, particularly for SSRT and proactive delaying, suggests that inferences about the neural mechanisms underlying inhibitory control do not generalize between group and individual levels. This nonergodic pattern highlights the importance of considering both levels of analysis to more fully understand the neurocognitive dynamics of inhibitory control.

### Network-level visualization of between- and within-subjects brain-behavior associations

To elucidate the patterns of brain-behavior relationships across different functional brain networks, we visualized the between-subjects and within-subjects associations for SSRT, probability of proactivity, and proactive delaying. We used the Shirer network atlas^48^ for our primary analysis as it includes the basal ganglia, a subcortical system important for the implementation of inhibitory control^24,49^.

*SSRT* (**Figure 4a**).

**Figure 4.**
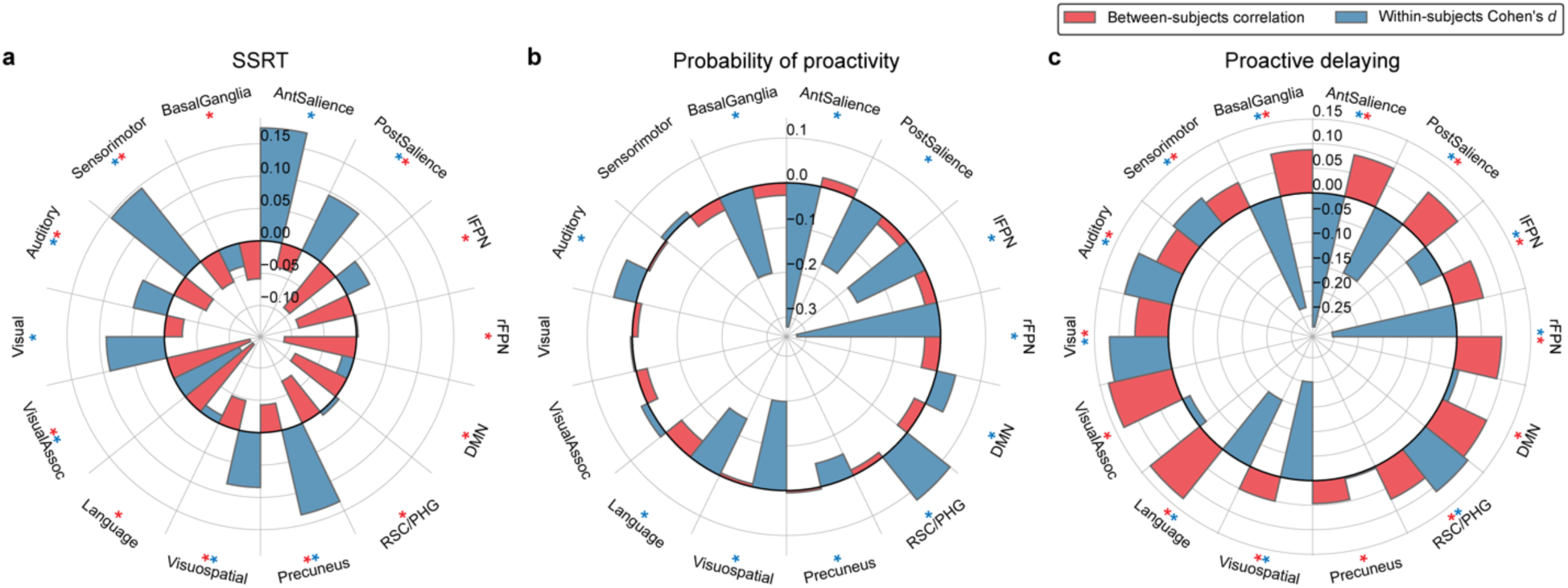
Network-level comparison of between-subjects and within-subjects brain- behavior associations in inhibitory control. The between-subjects analysis correlated subject- average brain activation (correct stop versus correct go activation) and subject-average cognitive model parameters: SSRT **(a)**, probability of proactivity **(b)**, and proactive delaying **(c)**. The within-subjects analysis regressed trial-by-trial brain activity on trial-by-trial cognitive model parameters: SSRT **(a)**, probability of proactivity **(b)**, and proactive delaying **(c)**. Effect sizes are shown for both analyses (between subjects: Pearson *r*; within subjects: Cohen’s *d*). Statistical significance is indicated by colored asterisks: red for between subjects (*P*_FDR_ < 0.01) and blue for within subjects (*P*_FDR_ < 0.01). Networks are based on the Shirer parcellation^48^. Differences in the existence and direction of associations between between-subjects and within-subjects analyses were observed across multiple brain networks, including the anterior and posterior salience, left and right frontoparietal, and default mode networks. AntSalience: anterior salience network; PostSalience: posterior salience network; lFPN: left frontoparietal network; rFPN: right frontoparietal network; DMN: default mode network; RSC/PHG: retrosplenial cortex / parahippocampal gyrus network; Precuneus: precuneus network; Visuospatial: visuospatial network; Language: language network; VisualAssoc: visual association network; Visual: visual network; Auditory: auditory network; Sensorimotor: sensorimotor network; BasalGanglia: basal ganglia network.

Between-subjects analysis showed consistent negative correlations in most networks, with effects in the posterior salience, frontoparietal, and default mode networks (all *P*_FDR_ < 0.01). In contrast, within-subjects analysis revealed mixed negative and positive associations, with opposite (positive) associations in the posterior salience, precuneus, visuospatial, auditory, and sensorimotor networks (all *P*_FDR_ < 0.01). The reversal of association directions underscores the nonergodic nature of SSRT-related brain activity.

#### Probability of proactivity (**Figure 4b**)

Between-subjects analysis demonstrated no significantly nonzero correlations in the networks (all *P*_FDR_ ≥ 0.05). However, within-subjects analysis unveiled a more complex pattern. Negative associations were observed in the salience and frontoparietal networks, while positive associations were seen in the default mode network (all *P*_FDR_ < 0.01). This disparity highlights the importance of examining within-subjects dynamics for proactivity.

#### Proactive delaying (**Figure 4c**)

Between-subjects analysis revealed positive correlations in all the networks (all *P*_FDR_ < 0.01). Yet, within-subjects analysis showed predominantly negative associations, notably in the salience and frontoparietal networks (all *P*_FDR_ < 0.01). The retrosplenial cortex and parahippocampal gyrus nodes of the ventral default mode network showed a positive within-subjects associations (*P*_FDR_ < 0.01).

These network-level visualizations emphasize the divergent patterns between group-level and individual-level associations across different functional brain networks. The consistent reversals observed, for both reactive and proactive control measures, reinforce that brain-behavior relationships in inhibitory control are nonergodic. These findings indicate that both between-subjects and within-subjects perspectives are needed to comprehensively understand the neural dynamics underlying inhibitory control.

### Stability of within-subjects brain-behavior associations

To assess the stability of our within-subjects findings, we performed bootstrap resampling at varying sample sizes (**Figure 5**). Within-subjects associations between brain activity and the model parameters (SSRT, probability of proactivity, and proactive delaying) were stable, even in modest sample sizes. In each of 5 independent sets of brain areas, resampled results showed high similarity with results in the full sample. Key findings were consistently observed in samples as small as 25 subjects, with some effects requiring larger samples to emerge reliably. This stability suggests the validity and potential generalizability of our within-subjects approach to understanding neurocognitive mechanisms of inhibitory control. A detailed description of these results is in the Supplementary Materials.

**Figure 5.**
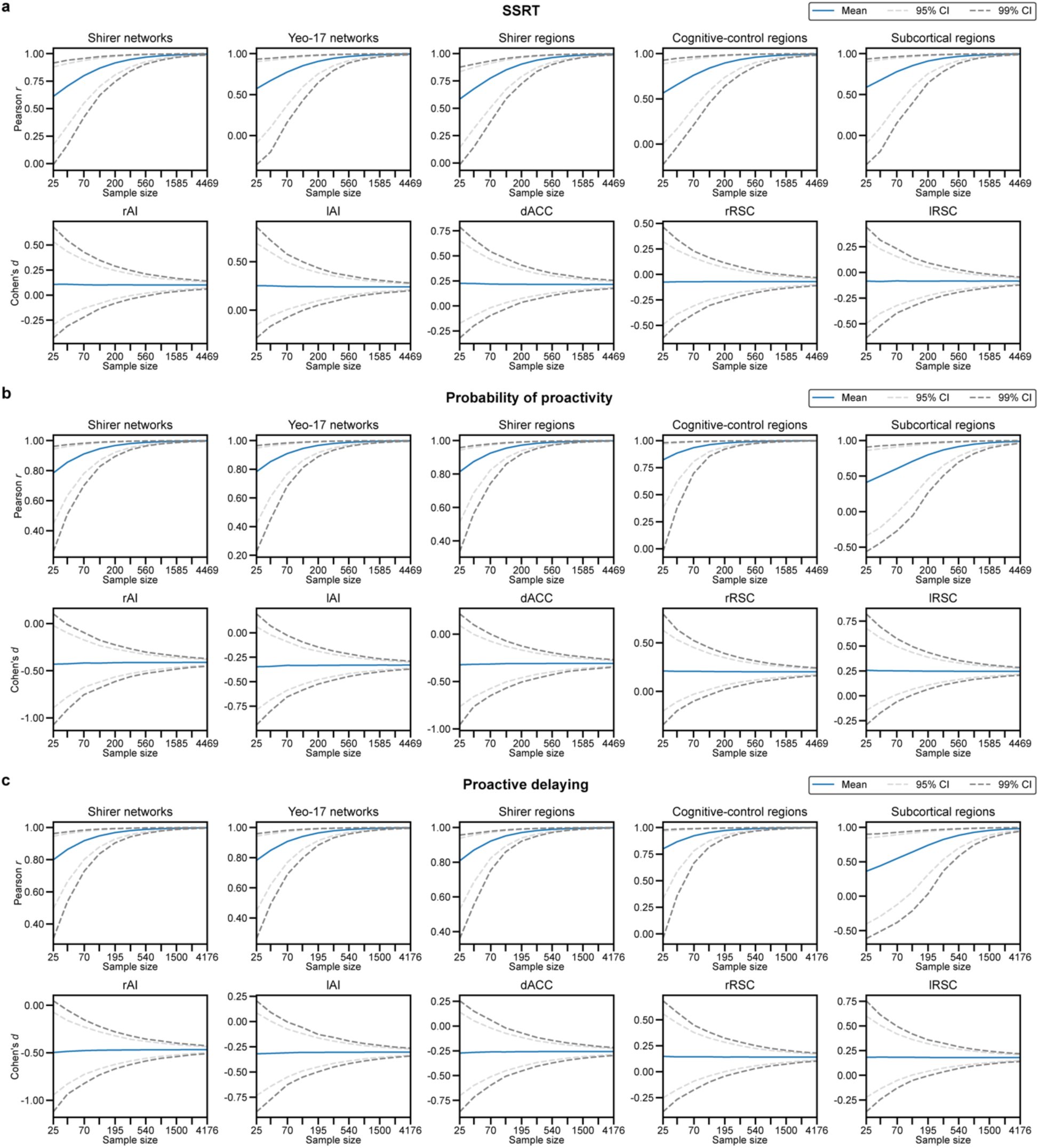
Stability analysis of within-subjects associations. **a-c**, Stability plots for SSRT **(a)**, probability of proactivity **(b)**, and proactive delaying **(c)**. In each panel, the top plots show the distributions of correlation between the effect in resamples and the effect in the full sample, while the bottom plots show the distributions of the effect in selected regions. For each plot, 10,000 resamples of *n* subjects were drawn with replacement for each sample size *n*, and Cohen’s *d* was calculated for each resample. For top panel plots, correlations were then computed for each resample over the areas belonging to the set of brain areas. The dashed lines depict 95% and 99% bootstrap confidence intervals. Within-subjects associations demonstrated stability for all 3 cognitive model parameters across 5 different collections of brain areas and in 5 regions of interest, even at modest sample sizes. This stability supports the reliability of brain-behavior associations in inhibitory control processes.

### Robustness of nonergodicity to analytical choices

To test the robustness of our findings of nonergodicity, we conducted several control analyses comparing between- and within-subjects brain-behavior relationships. We tested alternative between-subjects analyses using different measures of brain activation and explored whether the observed nonergodicity was specific to the latent model parameters or could also be seen using reaction time on go trials, a directly observed behavioral measure. Nonergodicity persisted for all approaches to between-subjects analysis of both model-derived parameters (**Supplementary Figure S2**) and the observed behavioral measure (**Supplementary Figure S3**). Across all analytical choices, we observed divergent patterns of between-subjects and within-subjects associations. A detailed description of these control analyses and their results is in the Supplementary Materials. Our results strongly suggest that the neurocognitive dynamics of inhibitory control in children are fundamentally nonergodic, with implications for how we interpret findings from traditional group-level associations.

### Representational similarity analysis reveals dissociated reactive and proactive representations

Building on our findings of nonergodic brain-behavior relationships, we sought to deepen our understanding of how the brain implements proactive and reactive control at the individual level. While the preceding results demonstrate the importance of within-subjects analyses, they leave unresolved the question of how representations of reactive and proactive processes relate to each other within brain networks. To investigate this, we used representational similarity analysis to examine the overlap between brain representations of reactivity (SSRT) and proactivity (probability of proactivity and proactive delaying) within individuals. For each subject and each brain network, we computed pairwise Pearson correlations between the subject’s brain maps of SSRT and probability of proactivity, SSRT and proactive delaying, and probability of proactivity and proactive delaying, over the voxels in each network (**Figure 6a**).

**Figure 6.**
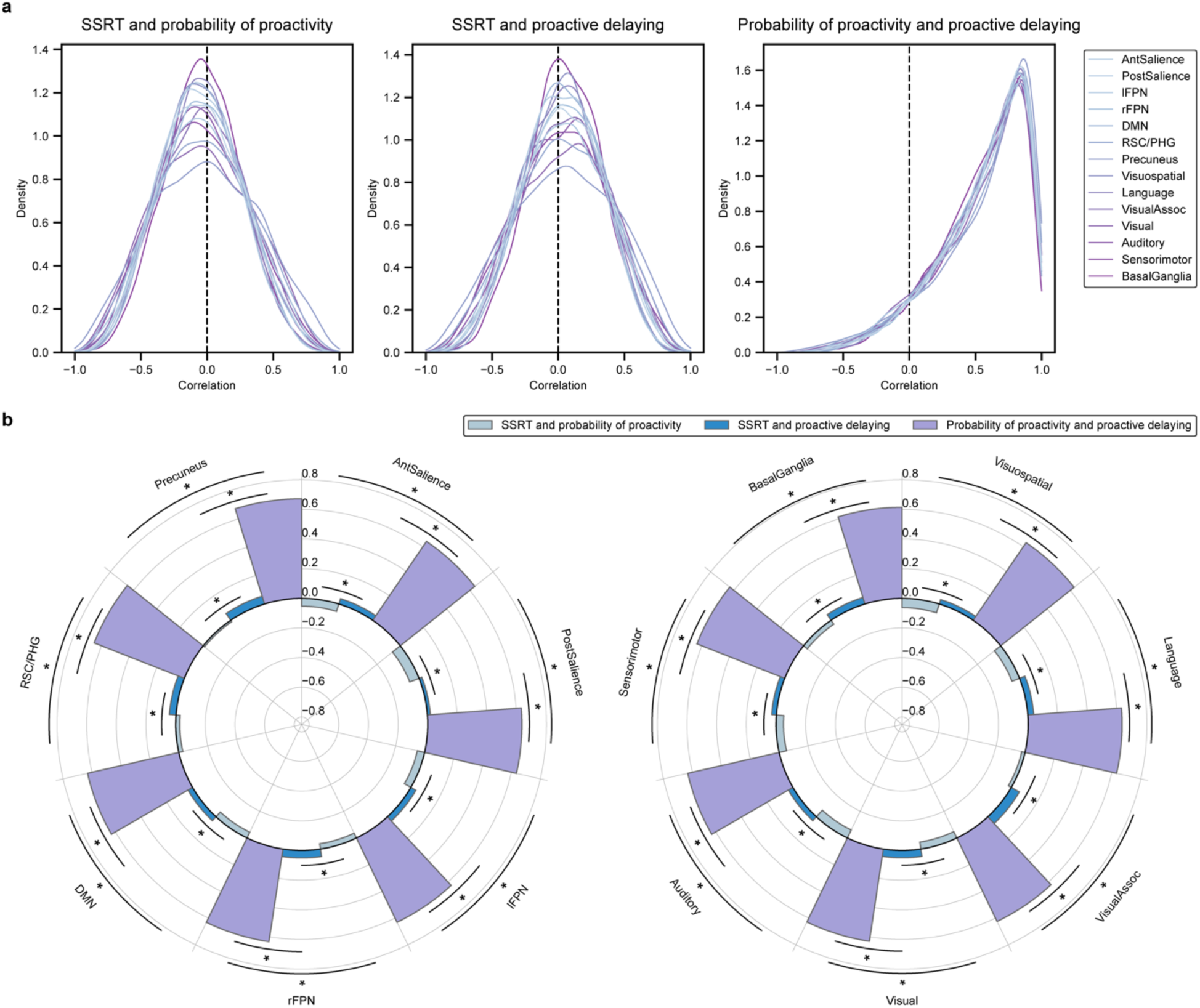
Dissociated brain representations of reactive and proactive control processes. For each subject, correlations were computed between pairs of within-subjects brain maps for the model parameters (SSRT, probability of proactivity, and proactive delaying) over each brain network. **a,** Distributions of correlations over subjects. Kernel density estimation was performed. **b,** Average correlations over subjects. The height of the bars is the median. Statistical significance of difference in medians is indicated by asterisks (* *P*_FDR_ < 0.01). Reactive (SSRT) and proactive (probability of proactivity, proactive delaying) control processes showed dissociated representations across all brain networks. This dissociation suggests that the brain employs distinct neural resources for reactive and proactive aspects of inhibitory control. In contrast, the two proactive measures showed high similarity, validating their representation of related cognitive processes.

SSRT showed low similarity with both proactive measures (probability of proactivity and proactive delaying) in all networks, with median correlations ranging from -0.07 to 0.06. In contrast, the 2 proactive measures (probability of proactivity and proactive delaying) exhibited high similarity in all networks, with median correlations ranging from 0.61 to 0.67. In each network, the 3 similarity measures (SSRT and probability of proactivity, SSRT and proactive delaying, and probability of proactivity and proactive delaying) were significantly different from each other in their median values (**Figure 6b**) (all *P*_FDR_ < 0.01). These findings suggest that representations of reactivity and proactivity are largely dissociated. This dissociation persists across multiple brain networks, indicating a fundamental separation in how the brain encodes reactive and proactive control processes.

### Adaptive regulation of inhibitory control associated with distinct within-subjects results over subgroups

To understand how our within-subjects associations related to between-subjects variation in cognitive and task strategies, we examined the within-subjects results between subgroups showing adaptive and maladaptive regulation of reactive and proactive behaviors. These subgroups were identified using PRAD model parameters γ_1_ (governing regulation of stopping expectancy, and hence SSRT over trials) and θ_1_ (governing regulation of probability of proactivity over trials, in response to performance monitoring).

First, the cognitive model infers for each subject γ_1_, which determines whether subjects adaptively (γ_1_ > 0) or maladaptively (γ_1_ < 0) regulate reactivity. Adaptive (maladaptive) regulation of reactivity involves increasing (decreasing) expectancy of a stop trial as the number of successive go trials increases. Since expectancy of a stop trial is one of several determinants of SSRT, γ_1_ influences SSRT variation through time. Thus, we examined within-subjects associations between SSRT and brain activity separately among subjects with γ_1_ < 0 (*N* = 2513) and γ_1_ > 0 (*N* = 1956) (**Figure 7a**). We found differences in the distributions of within-subjects SSRT associations between the γ_1_ subgroups in all networks examined (all *P*_FDR_ < 0.01). The maladaptive regulation group had a larger (more positive) effect size in every network. In fact, across most networks, SSRT exhibited opposite associations with brain activity in the two subgroups. For example, in the frontoparietal and default mode networks, SSRT displayed a positive association among subjects with γ_1_ < 0 and a negative association among subjects with γ_1_ > 0 (all *P*_FDR_ < 0.01). Moreover, this analysis revealed that some of the effects in the full sample were driven by subjects belonging to one of the subgroups. For example, in the anterior salience network, brain activity’s positive association with SSRT in the full sample was driven by γ_1_ < 0 subjects; among γ_1_ < 0, the effect in the anterior salience had a Cohen’s *d* of ∼0.3, twice that of the effect in the full sample, while among γ_1_ > 0, there was no significant effect at all.

**Figure 7.**
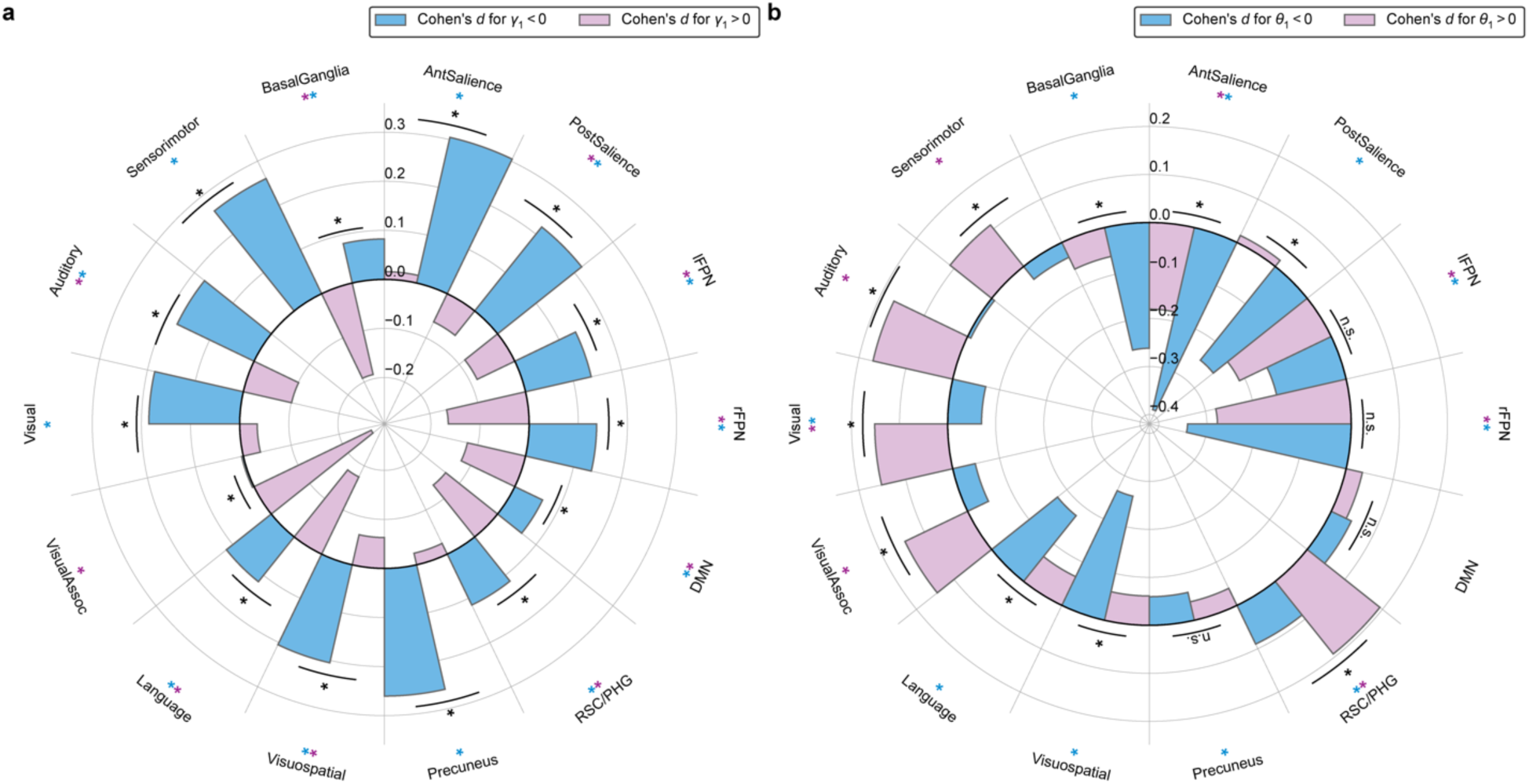
Distinct brain-behavior associations for adaptive and maladaptive regulators of inhibitory control. **a**, Within-subjects associations of SSRT with brain activity between γ_1_ subgroups. We identified two distinct subgroups of subjects with opposite profiles of attentional modulation. Subjects showed either maladaptive regulation (γ_1_ < 0, *N* = 2513) or adaptative regulation (γ_1_ > 0, *N* = 1956) of their expectancy of stopping over time. The two groups showed differences in within-subjects associations between SSRT and brain activity in various networks. This demonstrates that individual differences in attentional dynamics play a role in shaping the relationship between neural activity and inhibitory control processes. A colored asterisk indicates that a network or region’s associations were nonzero among subjects with γ_1_ < 0 (blue * *P*_FDR_ < 0.01) or among subjects with γ_1_ > 0 (purple * *P*_FDR_ < 0.01). **b,** Within-subjects associations of probability of proactivity with brain activity between θ_1_ subgroups. We identified two distinct subgroups of subjects with opposite profiles of performance monitoring. Subjects showed either adaptive regulation (θ_1_ < 0, *N* = 3054) or maladaptive regulation (θ_1_ > 0, *N* = 1415) of their proactivity over time. The two groups showed differences in within-subjects associations between probability of proactivity and brain activity in various networks. This demonstrates that the neural correlates of proactive control are influenced by an individual’s strategy for adjusting proactivity in response to task outcomes. A colored asterisk indicates that a network or region’s associations were nonzero among subjects with θ_1_ < 0 (blue * *P*_FDR_ < 0.01) or among subjects with θ_1_ > 0 (purple * *P*_FDR_ < 0.01). For both panels, a black asterisk indicates that associations had different distributions between the two subgroups (* *P*_FDR_ < 0.01, n.s. *P*_FDR_ ≥ 0.01).

Second, the cognitive model infers for each subject θ_1_, which determines whether subjects adaptively (θ_1_ < 0) or maladaptively (θ_1_ > 0) regulate proactivity. Adaptive (maladaptive) regulation of proactivity involves increasing (decreasing) the probability of proactivity following a failed stop trial and decreasing (increasing) the probability of proactivity following a no-response go-trial. Therefore, θ_1_ influences the probability of proactivity’s variation through time. Thus, we examined the probability of proactivity’s within-subjects associations with brain activity separately among subjects with θ_1_ < 0 (*N* = 3054) and θ_1_ > 0 (*N* = 1415) (**Figure 7b**). We found consistent differences in the probability of proactivity’s within-subjects associations between these subgroups. Generally, subjects with adaptive regulation (θ_1_ < 0) had more negative associations between probability of proactivity and brain activity. This negative coupling between proactivity and brain activity was pronounced within the adaptive regulation group in the anterior salience network (Cohen’s *d* of ∼0.4). This analysis also clarified how adaptive and maladaptive regulation of proactivity contributed to the effects observed in the full sample. The within-subjects positive association between probability of proactivity and activation in the retrosplenial cortex and parahippocampal gyrus network was observed in both θ_1_ subgroups (both *P*_FDR_ < 0.01), but it was twice as large among subjects who maladaptively regulated proactivity.

These results demonstrate that population subgroups related to adaptive and maladaptive regulation of inhibitory control demonstrate different, and even opposite, within-subjects brain-behavior associations.

### Brain networks exhibit varying degrees of nonergodicity and a hierarchical organization by nonergodicity

To further understand the distribution of nonergodicity across the brain, we quantified and compared nonergodicity levels in different brain networks. We defined a nonergodicity measure as the fraction of subjects for whom within-subjects brain-behavior association showed an opposite sign to the between-subjects association. Values above 0.5 indicate higher nonergodicity, while values below 0.5 suggest more ergodic behavior.

Our analysis revealed substantial variation in nonergodicity across brain networks (**Figure 8a**). Notably, the anterior salience network consistently demonstrated the highest level of nonergodicity for all three cognitive model parameters (SSRT, probability of proactivity, and proactive delaying). The measures of network nonergodicity for associations with the probability of proactivity showed wide confidence intervals, reflecting the weak between-subjects associations between this parameter and brain activation.

**Figure 8.**
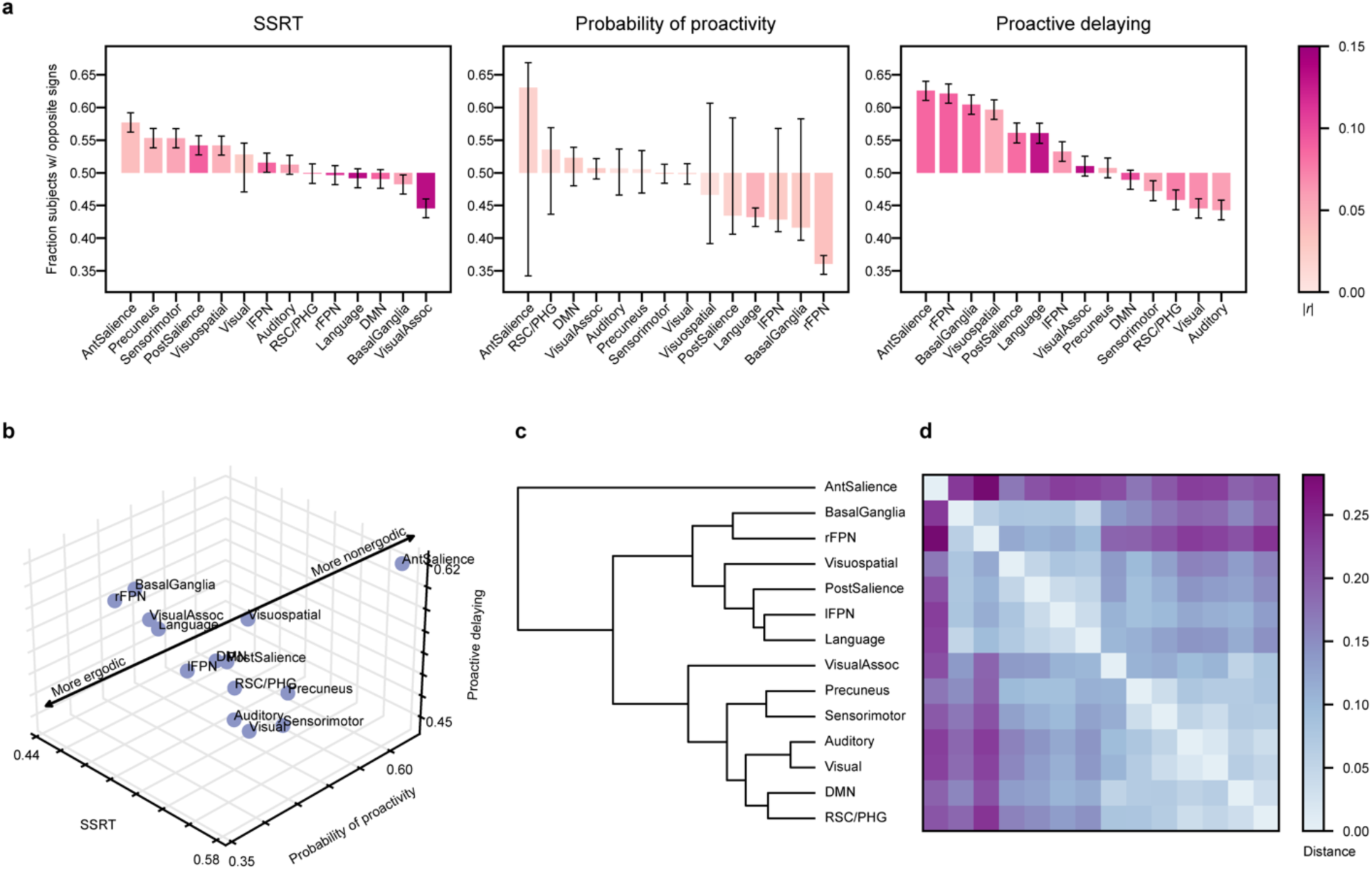
Nonergodicity across brain networks. **a**, Degree of nonergodicity of brain networks. Bar plots show the extent of nonergodicity for each brain network across three cognitive model parameters (SSRT, probability of proactivity, proactive delaying). The height of the bars is the mean fraction of subjects showing opposite-sign associations compared to between-subjects results. The bars are shaded by the magnitude of the between-subjects correlations. Error bars show 95% bootstrap confidence intervals. **b,** 3-dimensional embedding of each network. Networks are represented as points based on their nonergodicity measures for the three parameters. **c-d,** Hierarchical clustering of networks based on their nonergodicity profiles. **c,** Dendrogram of clustering. **d,** Euclidean distances between network embeddings. Brain networks exhibited varying degrees of nonergodicity. The anterior salience network showed the highest level of nonergodicity and a unique profile of nonergodicity distinct from that of all other networks.

To understand how networks related to each other in terms of nonergodicity, we performed hierarchical clustering on the joint nonergodicity measures of the networks with respect to all three cognitive model parameters (**Figure 8b-d**). This analysis revealed a hierarchical organization of brain networks based on their nonergodicity profiles. The anterior salience network emerged as the most dissimilar, forming its own cluster separate from all other networks. Additionally, the default mode network and retrosplenial cortex and parahippocampal gyrus clustered together, suggesting similarities in their nonergodic behavior.

These findings demonstrate that brain networks exhibit varying levels of nonergodicity and reveal a hierarchical organization of brain networks based on their nonergodic properties. This organization suggests a new perspective on the functional architecture of the brain and how it relates to cognitive control processes.

## Discussion

A fundamental question we examined in this study is whether between- and within-subjects brain-behavior associations yield convergent findings in understanding neurocognitive processes underlying cognitive control. Divergence between these levels of analysis would provide evidence for nonergodicity, a phenomenon where inferences drawn from population-level data do not accurately represent individual-level processes. To investigate this, we leveraged a large community sample of children from the ABCD study and employed a dynamic computational model to elucidate the potentially nonergodic nature of neurocognitive dynamics underlying cognitive control. We combined task fMRI data with a hierarchical Bayesian model of cognitive dynamics to examine brain-behavior relationships at both between-subjects and within-subjects levels. Our findings reveal pervasive nonergodic patterns in brain-behavior relationships underlying cognitive control, with striking divergences between population-level and individual-level analyses. These results underscore the need to distinguish between-subjects and within-subjects inferences about the brain—a new paradigm we term “nonergodicity neuroscience”.

At the between-subjects level, inhibitory control activations in key cognitive control networks were negatively correlated with subject-average SSRTs, aligning with previous studies^24,35–37^. This suggests that individuals with better inhibitory control (lower SSRTs) exhibit greater activation in regions associated with cognitive control. However, within-subjects analysis revealed a markedly different pattern: brain activity in some of these same networks was positively associated with trial-level SSRTs and negatively associated with trial-level proactive control. Thus, within-subjects associations were largely dissonant with the between-subjects findings, revealing the nonergodic nature of inhibitory control processes. Specifically, the opposing patterns demonstrated Simpson’s paradox, a type of nonergodicity frequently observed in behavioral studies, where relationships at the group level are changed at the individual level^5,12,13,50^. This divergence underscores the importance of considering both group-level and individual-level associations in cognitive neuroscience research and highlights the complex, dynamic nature of inhibitory control processes.

Furthermore, representational similarity analysis uncovered distinct neural representations for reactive and proactive cognitive processes. Across all examined brain networks, representations of reactive control showed low similarity with both measures of proactive control, while the two proactive measures showed high similarity with each other. This dissociation suggests that reactive and proactive aspects of inhibitory control rely on distinct neural resources, potentially allowing for independent modulation and development of these strategies.

### Modeling trial-level responses at the single-subject level

We used PRAD, a novel hierarchical Bayesian model of proactive and reactive control which represents a significant advance in the assessment of inhibitory control, surpassing the capabilities of conventional race models^44^. Unlike traditional approaches, which provide subject- aggregate SSRT estimates as an index of inhibitory control, PRAD estimates SSRT at the level of each individual trial. Briefly, PRAD models trial-specific SSRT estimates that account for both proactive control strategies and dynamic attentional modulation effects, providing dynamic measures of latent inhibitory control processes^44^. This feature is critical as it allows for precise, trial-specific inferences rather than broad generalizations across the entire task. Importantly, the trial-level model infers additional trial-level measures to the SSRT, such as the probability of proactive cognitive states and the length of proactive delaying of responses.

Trial-level granularity facilitated a more comprehensive examination of how neural responses are modulated across individual trials. Specifically, this model allowed us to identify brain areas that tracked neural activity in response to trial-specific SSRT estimated at each stop trial. Moreover, it enabled tracking ongoing neural dynamics associated with temporal fluctuations in proactive control, providing insights into the brain systems that support a key component of cognitive control.

By employing trial-level analyses, we could dissect proactive and reactive control processes to examine how they fluctuate over time within each individual. This approach deepens our understanding of the mechanisms underlying inhibitory control and enabled us to determine whether assumptions of brain-behavior ergodicity are justified.

### Nonergodic brain-behavior associations reveal novel mechanisms of cognitive control

Nonergodicity, a concept originating from physics, describes situations where ensemble averages and time averages do not converge^1^. In the behavioral and life sciences, nonergodicity manifests when measures such as the mean, variance, or covariance differ between population-level analyses (between subjects) and individual-level analyses over time (within subjects)^5,10,11,16^. Our study focused on comparing the covariance of brain activity and behavior between these two levels of analysis, considering divergent associations as evidence of nonergodicity and Simpson’s paradox^12,42,50,51^.

Our findings reveal pervasive nonergodic patterns in brain-behavior relationships across multiple measures of inhibitory control, encompassing both proactive and reactive control processes. Proactive control, involving anticipation and preparation for stopping, and reactive control, involving the actual implementation of response inhibition, showed markedly different patterns at the within-subjects level compared to the between-subjects level.

We found a positive association within subjects between trial-level SSRTs and brain activity in the anterior and posterior salience networks, which suggests that longer SSRTs, indicating poorer reactive control, are associated with greater neural effort or engagement. This may reflect compensatory mechanisms or the increased demand for cognitive resources when individuals struggle to inhibit their responses. In contrast, between subjects, we observed no significant association and a negative association, respectively, in the anterior and posterior salience networks between SSRT and brain activity.

Within-subjects analysis revealed a negative association between trial-level engagement of proactive control and brain activity in the frontoparietal, salience, and subcortical systems. This is consistent with greater proactive control being associated with a lower need to engage cognitive control networks that facilitate reactive control, to implement successful response inhibition. This finding also aligns with recent theoretical frameworks proposing that proactive control modulates reactive control via preparatory processes^52^. In contrast, we observed a positive association between trial-level proactive control and default mode network activity. This included the posterior medial cortex and the ventromedial prefrontal cortex, the two core cortical nodes that anchor the default mode network^53^. This may reflect internally oriented processing that supports proactive regulation. Between-subjects analysis failed to capture these dynamic relationships, showing no association between one measure of proactive control and brain activations.

Our findings highlight the importance of considering within-subjects variability and dynamics when studying the neural mechanisms of cognitive control. Conventional between-subjects analyses, which assume ergodicity, may not capture the complex and dynamic nature of proactive and reactive control processes as they unfold within individuals over time during development and in psychopathology. Nonergodicity neuroscience approaches using the methodologies advanced in this study may be crucial for understanding neurocognitive functions at the individual subject level.

### Robustness of nonergodicity findings and stability of within-subjects brain-behavior associations

Leveraging the large-scale ABCD dataset, we addressed the critical challenge of replicability in human neuroscience^54,55^. Our analyses revealed that within-subjects associations were stable and reliably detectable, even in sample sizes typical of cognitive neuroscience studies. Bootstrap resampling analyses showed the reliability of key findings across different sample sizes. For instance, the association between proactivity measures and right anterior insula suppression was consistently observed in over 95% of samples, even with sample sizes of *N* = 25. Moreover, our findings of nonergodicity were robust to various analytical strategies. When comparing the results of various between- and within-subjects approaches, there were variations in the details of brain-behavior inferences, but nonergodic dissociations persisted across every approach to brain-behavior association, including the use of observed dynamic measures (go reaction time) rather than model-based latent dynamic measures.

By demonstrating that these patterns are robust and detectable even in modest sample sizes, our study provides a foundation for future research into nonergodicity in brain function. It also suggests that meaningful insights into neurocognitive mechanisms can be gained from studies with more typical sample sizes, although larger samples provide greater precision and the ability to detect subtler effects. The stability and robustness of our findings suggests their applicability to diverse research and clinical contexts, including understanding cognitive processes related to inhibitory control and studying psychiatric disorders.^56^

### Nonergodicity between brain networks: Implications for understanding cognitive control

Our analysis of the distribution of nonergodicity between brain networks has implications for understanding the neural mechanisms underlying inhibitory control as well as for cognitive neuroscience research broadly. The consistent finding of high nonergodicity in the anterior salience network across all cognitive model parameters is particularly intriguing. The salience network is a key brain network, known for its role in detecting behaviorally relevant stimuli and coordinating brain network dynamics^57,58^. It is noteworthy that this core network, which is of great interest in the study of cognition^23–26,59^ and psychopathology^60–62^, appears to exhibit the most pronounced disconnect between group-level and individual-level inferences. Moreover, the hierarchical clustering of networks based on nonergodicity profiles provides a fresh perspective on brain organization. The distinct clustering of the anterior salience network and the default mode network suggest that nonergodicity may be an important factor in understanding functional brain architecture.

These findings have several significant implications for cognitive neuroscience research and practice. Methodologically, our results underscore the importance of complementing group-level analyses with individual-level investigations. The high degree of nonergodicity observed, particularly in key networks involved in cognitive control, suggests that solely relying on group-level analyses may lead to incomplete or misleading conclusions about brain-behavior relationships. From the perspective of individual differences, the varying levels of nonergodicity across networks highlight the importance of considering individual variability in brain function. This may be particularly relevant for understanding individual differences in inhibitory control abilities and for developing personalized interventions for disorders characterized by impaired inhibitory control. Theoretically, the observed nonergodicity challenges simplistic models of brain function and calls for quantitatively rigorous theories that can account for the complex, context-dependent nature of brain-behavior relationships. This may require a shift toward more dynamic, process-oriented models of cognition and brain function.

### Distinct neural representations for reactive and proactive cognitive processes

To further elucidate the neural architecture underlying inhibitory control, we employed representational similarity analysis, a powerful method for investigating the informational content of brain activity patterns^63^. This approach allows us to compare the similarity of neural representations across different cognitive processes, providing insights into how the brain organizes and processes information^64^. In our study, we used representational similarity analysis to examine the overlap between brain representations of reactivity (SSRT) and proactivity (probability of proactivity and proactive delaying) within individuals. By comparing representational patterns across different cognitive processes, we sought to determine whether reactive and proactive control rely on shared or distinct neural resources.

Our analysis revealed a striking dissociation between the neural representations of reactive and proactive control processes. Across all examined brain networks, we found low similarity between representations of SSRT and representations of each proactive measure (probability of proactivity and proactive delaying). In contrast, the two proactive measures showed high similarity with each other. This pattern suggests that reactive and proactive control processes are represented orthogonally in the brain. These measures of proactivity affecting the go process and reactivity affecting the stopping process are more explicitly defined based on naturally occurring dynamics as inferred by the PRAD model in the standard SST task, rather than being based on modified versions of the standard task which may attempt to experimentally induce proactive and reactive aspects of inhibitory control on different trials^29,65–67^. Theoretically, our findings challenge simplistic models of inhibitory control and suggest that reactive and proactive processes, while both contributing to inhibitory control, are implemented through distinct neural mechanisms^46,68^.

Given that our study focused on children, the clear separation of reactive and proactive representations may reflect a developmental stage in the organization of cognitive control processes. The separation we found may allow for independent development of reactive and proactive strategies, potentially explaining individual differences in inhibitory control abilities^46^. Future studies could investigate whether this orthogonality persists or changes with age^69^.

### Attentional modulation and performance monitoring associated with distinct brain-behavior associations

A separate behavioral investigation of our hierarchical Bayesian model highlighted the significance of adaptive regulation of reactive and proactive control in shaping within-subjects variability in SSRT and stop failure rates^44^. Two model parameters are decisive in controlling these dynamics: γ_1_ represents individual differences in sustained attention and regulates the trial-level expectancy of stopping, and θ1 represents individual differences in performance monitoring and regulates the trial-level proclivity for proactive control. These findings point to the importance of considering individual differences in attentional modulation and performance monitoring systems when studying inhibitory control.

Building on these results, we investigated brain-behavior associations between subjects who differed on these traits. We used γ_1_ and θ_1_ (separately) as a basis for creating subgroups within our sample. By dividing our participants into subgroups, we could examine how individual differences in attentional regulation and performance monitoring correlate with the relationship between neural activity and cognitive processes within subjects. Subjects stratified based on whether they adaptively or maladaptively regulated stopping expectancy showed distinct within-subjects associations between trial-level SSRTs and brain activity, with different distributions of associations between the subgroups in every network and opposite associations in most networks. Among all subjects, we observed that anterior salience network activation accompanied poorer reactive control at the trial level, but examining these results by γ_1_ subgroup revealed that this association only held for subjects who maladaptively regulated reactivity. Similarly, subjects divided by whether they adaptively or maladaptively regulated proactivity showed different within-subjects associations for a measure of proactivity in most networks. With greater trial-level proactivity, the maladaptive regulation group showed weaker suppression of the anterior salience network and stronger activation of the retrosplenial cortex and parahippocampal gyrus network, which anchors the ventral posterior aspects of the default mode network.

Collectively, the findings reveal that groups characterized by adaptive and maladaptive regulation of reactivity and proactivity display notably different patterns of within-subjects associations between brain activity and model parameters. These distinctions hint that individuals’ distinct cognitive strategies or profiles relate to the implementation of proactive and reactive control processes in the brain. This variation highlights the personalized nature of cognitive function and stresses the importance of considering individual differences in the neural mechanisms of inhibitory control. Identifying heterogeneity based on cognitive model parameters provides an interpretable approach for studying individual differences in inhibitory control and their neural correlates. This approach moves beyond simple between-subjects comparisons and allows for a theory- and mechanism-driven investigation of the heterogeneity in brain-behavior relationships.

### Conclusions

Our study provides evidence for nonergodicity in the neurocognitive processes underlying inhibitory control using a large, community-representative sample of children from the ABCD study. By combining task fMRI data with a dynamic cognitive model, we found that within-subjects associations between brain activity and model parameters differed from between-subjects associations, challenging the assumption of ergodicity in cognitive neuroscience research. The findings demonstrate divergent group-level and individual-level brain-behavior associations, reveal dissociated proactive and reactive control systems, and identify meaningful individual differences in these control processes. Crucially, the study establishes the stability and robustness of within-subjects measures, laying a foundation for characterizing nonergodic processes. The work highlights the value of large, heterogeneous samples, dynamic computational models, and analysis of within-subjects variability for studying nonergodic phenomena. More broadly, it suggests a paradigm shift away from untested ergodic assumptions toward nonergodicity neuroscience. Appreciating the nonergodic nature of neurocognitive processes may be essential for advancing our understanding of cognition in health and disease, as well as of artificial neural networks.

## Acknowledgements.

This work was supported by National Institutes of Health MH121069 (V.M.), MH124816 (W.C.), National Science Foundation No. 2024856 (V.M.), and the Stanford Maternal and Child Health Research Institute (W.C., P.M.).

## Contributions

Study design: P.K.M., N.K.B., Z.G., W.C., V.M.; Analysis: P.K.M., N.K.B., Z.G.; Interpreting results: P.K.M., N.K.B., Z.G., W.C., V.M.; Writing: P.K.M., N.K.B., V.M.; Editing: P.K.M., N.K.B., Z.G., W.C., V.M.; Supervision: P.K.M., V.M.

## Data availability

Data used in this study were from the ABCD study (https://abcdstudy.org/), held in the National Institute of Mental Health Data Archive. These data are publicly shared with eligible researchers.

## Code availability

All code used in this study will be made available upon publication (https://github.com/scsnl).

## Methods

### Inclusion criteria

Data were from the baseline visit of the ABCD study^18^ (Collection #2573), *N* = 11817. Subjects were excluded if they did not meet each of the following criteria: meet the ABCD study’s SST task-fMRI inclusion recommendations (in abcd_imgincl01.txt, imgincl_sst_include==1; *N* = 3546 excluded); have 2 SST fMRI runs of good quality (in mriqcrp20301.txt, iqc_sst_total_ser==iqc_sst_good_ser==2; *N* = 677 excluded); are successfully fit with the cognitive model of the SST (*N* = 562 excluded); have 2 SST fMRI runs in the release 4.0 minimally processed data (*N* = 16 excluded); have enough volumes acquired to cover the SST experiment (the last SST trial must have happened no more than 2 seconds after the final volume was acquired; *N* = 16 excluded); have mean framewise displacement of less than 0.5 mm for both runs (calculated using the method of ref. ^70^; *N* = 1986 excluded); have release 4.0 minimally processed events.tsv files of shape (181,3) for both runs (*N* = 7 excluded); and have consistent release 4.0 behavioral data (in release 4.0, for some subjects, the “sst.csv” files from ABCD Task fMRI SST Trial Level Behavior, abcd_sst_tlb01, disagreed with the minimally processed “events.tsv” files; for example, one trial might be labeled a go trial by one file and a stop trial by the other; *N* = 102 excluded). Then, we excluded siblings by randomly keeping one member from each family (using the genetic_paired_subjectid variables from gen_y_pihat; *N* = 436 excluded) and excluded subjects without scanner serial number recorded (in mri_y_adm_info, missing mri_info_deviceserialnumber; *N* = 10 excluded). Applying these inclusion criteria left us with a sample of *N* = 4469. For analyses involving the proactive delaying, a further 293 subjects were excluded who had no trials with probability of proactivity greater than 0.5 during at least one run, and therefore, by definition, a proactive delaying of 0 for all trials of at least one run. For these subjects, we were unable to examine within-subjects relationships between proactive delaying and brain activity. To maintain comparability of the between- and within-subjects analyses, we also excluded these subjects from the between-subjects analyses involving proactive delaying. Thus, analyses involving the proactive delaying used a sample of *N* = 4176.

### Brain imaging

Imaging acquisition for the ABCD SST is detailed in other work.^71^ We used the minimally processed data from ABCD release 4.0 (Collection #2573), which included distortion correction and motion correction^72^. We then further processed the images using Nilearn and FSL FLIRT: (1) initial volumes were removed (Siemens: 8, Philips: 8, GE DV25: 5, GE DV26 and other GE versions: 16); (2) the mean image in the time dimension was computed using mean_img from the Nilearn image module; (3) the mean image was registered to an echo-planar imaging template in MNI152 space (SPM12’s toolbox/OldNorm/EPI.nii) using FSL FLIRT, and an affine of this transformation was obtained; (4) the time-series of images was spatially normalized to MNI152 space with the affine from the previous step using FSL FLIRT; and (5) the images were smoothed with a Gaussian filter with a full-width at half maximum of 6 mm using smooth_img from the Nilearn image module.

### Bayesian modeling of cognitive dynamics

The PRAD model^44^ incorporates latent dynamics that respond to endogenous and exogenous variables, with trait measures governing the interaction of such endogenous and exogenous variables with latent processes, giving rise to non-stationary dynamics. This allows the PRAD model to account for violations of context and stochastic independence. Overall, the PRAD model incorporates separate evidence accumulation (drift-diffusion) processes for the go and stop processes, similar to a canonical horse-race model^73^. However, in addition to typical drift-diffusion process parameters, PRAD includes additional individual trait-like and dynamic trial-level measures. The full PRAD model is specified below.

(a) The go process is modeled as a drift diffusion process, with a trial-invariant non-decision time (τ_*G*_) and initial directional bias (β_*G*_), but a trial-varying decision threshold (α_*G,t*_) and drift rate (δ_*G,t*_). The dynamic decision threshold

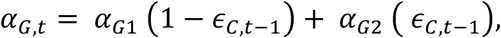

where (α_*G*1_, α_*G*2_) are distinct threshold levels and ϵ_*c,t*–1_ is an indicator of whether the left vs right choice on the previous trial was erroneous. Thus, the dynamic threshold implements a form of performance monitoring and varies between two levels based on the outcome of the previous trials, with α_*G*1_ being the default threshold and α_*G*2_ reflecting the threshold after post-error adjustments. The dynamic drift rate

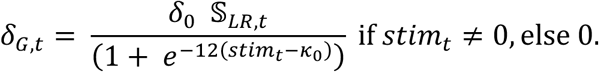

Here δ_0_ is a measure of the maximum drift rate for an individual, with the actual drift rate depending on the duration of the go stimulus (*stim_t_*) and an individual parameter κ_0_, which can be interpreted as the stimulus duration at which the drift rate is half the maximum. 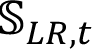 assumes values 1 or -1 depending on the direction of the go stimulus (left or right). PRAD allows for trial level changes to the drift rate, overcoming the issues with variable go stimulus durations highlighted in previous work^74^.

In addition, in the PRAD model, the onset of the go process may be deliberately delayed in anticipation of a stop signal. This dynamic adaptation is modeled by adding a further delay ω_*t*_ to the go process to reflect proactive delayed responding to the go stimulus, where

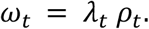

Here, λ_*t*_ reflects a trial-level belief updating process, based on the history of stop signal delays (SSD) encountered, and is an internal noisy estimate of the prospective anticipated SSD. The parameter μ (0 < μ < 1) reflects persistence in belief updating, with high persistence implying a lower decay rate of older SSDs encountered. Further, letting *SSD_t_* be the SSD and 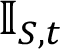 be a stop trial indicator,

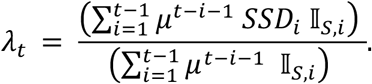

ρ_*t*_ is a binary variable representing cognitive state. ρ_*t*_ indicates the presence (ρ_*t*_ = 1) or absence (ρ_*t*_ = 0) of a proactive cognitive state on trial *t*. The proactive delayed responding is only initiated on proactive cognitive states. Proactive cognitive states are governed by a baseline proclivity for proactivity (θ_+_), and a performance-monitoring based modulation (θ_1_).

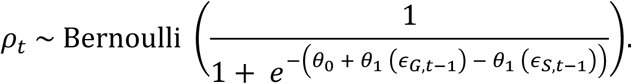

Here, ϵ_*G,t*−1_ is an indicator of a go-omission (incorrectly stopping on a go trial) on the previous trial, and ϵ_*S,t*−1_ is an indicator of a stopping error (not stopping on a stop trial). The PRAD model assumes that the correction in terms of increasing or decreasing the probability of a proactive cognitive state following these two types of trials will be in opposite directions. The sign of θ_1_ is an indicator of adaptivity or maladaptivity of the performance monitoring mechanism, and the absolute value of θ_1_ denotes the sensitivity of the state-switching mechanism to errors.

(b) The stop process is modeled as a drift diffusion process with a trial-invariant non-decision time (τ_*s*_), decision threshold (α_*s*_), and drift rate (δ_*s*_), but a trial-varying bias (β_5,’_). The stop process begins at the onset of the stop signal. The initial bias is

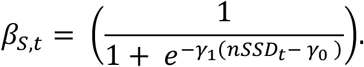

Here, *nSSD_t_* reflects the number of trials since a stop signal was last encountered. This reflects an attentional mechanism that modulates the stopping bias β_*s,t*_ (which varies from 0 to 1). Positive values of γ_1_ result in an increase in stopping bias as *nSSD_t_* increases and vice versa. Similarly, negative values of γ_1_ result in a decrease in stopping bias as *nSSD_t_* increases and vice versa. The absolute value of γ_1_ measures the sensitivity to attentional modulation. The γ_0_ parameter is a measure of the value of *nSSD_t_* when stopping bias is neutral (0.5).

Both the go and stop processes are implemented within a hierarchical Bayesian modeling framework in JAGS^75^, using the Wiener distribution^76^, which produces a joint distribution of the reaction times and the decision choice on each trial. The reaction times of the go process correspond to the reaction times for pressing the left or right buttons in response to the go stimulus. The reaction times of the stop process correspond to the SSRT. The stop process is only initiated on stop trials after the appearance of the stop stimulus (which appears after a delay corresponding to the SSD). The SSRT is not manifested as a behavioral action. Rather, if the SSRT, which is the duration of the stop process, plus the SSD on a stop trial is smaller than the go process reaction time, then the go action can be successfully inhibited (successful stop). The interaction of the basic go and stop processes can be influenced by the dynamics of the proactive delayed responding as well as the dynamics of the attentional modulation of reactive stopping.

The PRAD model enables obtaining the full posterior distributions of SSRT, proactive delay in responding of the go process, and the probability of proactive cognitive states at a trial level.

Markov Chain Monte Carlo was run using 4 chains with 30,000 samples each. For each chain, we had a burn-in of 10,000 samples, and a thinning factor of 10 was used to record 2000 of the remaining 20,000 samples, based on which final inferences were made. The following priors were used. δ_*G*_∼Uniform(0.0001,12). α_*G*1_∼Uniform(0.5,4). α_*G*2_∼Uniform(0.5,4). κ_+_∼Normal(0.1, precision = 0.3) truncated to (0,0.3). β_*G*_∼Beta(1,1) truncated to (0.0001,0.9999). δ_*s*_ = *p*/α_*s*_, where *p*∼Uniform(4,24) and α_*s*_∼Uniform(0.5,6). θ_+_∼Normal(0, precision = 0.3). θ_1_∼Normal(0, precision = 0.3). μ∼Beta(0.5,0.5) truncated to (0.8,0.9999). γ_1_∼Normal(0, precision = 0.3). γ_+_∼Uniform(0,12). τ_*G*_∼Uniform=0.0001, min(0.1, RT^∗^)@, where RT^∗^ is the reaction time after censoring for extremely small values. τ_*s*_∼Uniform(0.0001, τ_*G*_). For further details of the model, see ref. ^44^. We defined the 3 latent dynamic measures examined in this study as follows. SSRT was the posterior mean of the duration of the stop process on correct or incorrect stop trials; it was not defined for go trials or trials on which a participant responded before the stop signal was shown. Probability of proactivity was the posterior mean of ρ_*t*_, i.e., the posterior probability of ρ_*t*_= 1. Proactive delaying was the posterior mean of λ_*t*_ on trials with probability of proactivity greater than 0.5, otherwise 0. Probability of proactivity and proactive delaying were defined for all trials.

### Between- and within-subjects, general linear model analysis of fMRI

We fit general linear models to the fMRI BOLD recordings using Nilearn’s FirstLevelModel. Condition and parametric regressors were modeled as impulses, with a duration of 0, and convolved with the SPM software’s double gamma hemodynamic response function and the function’s time derivative. Before fitting, the BOLD signal was scaled to percent signal-change from the mean in the time dimension. An AR1 model was used to whiten the data and design matrices to account for temporal autocorrelation in the BOLD signal.

To determine between-subjects associations between subject-average brain activity and behavioral measures, for each subject and voxel, we fit the model

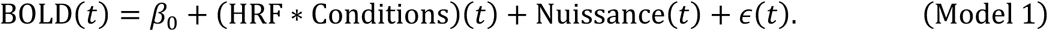

BOLD(*t*) is the BOLD signal of the voxel at time *t* (*t* ∈ {1, …, *T*} for *T* the total number of volumes acquired); (HRF ∗ Conditions)(*t*) is the value at *t* of the convolution with the hemodynamic response function HRF of condition regressor(s) Conditions; Nuissance(*t*) is the effect at *t* of nuisance regressors, which were 6 motion parameters (translational and rotational displacement along each of three axes) and 6 cosine basis functions (corresponding to high-pass filtering at 0.01 Hz); and ϵ(*t*) is the model’s error at *t*. We fit models with three sets of condition regressors:

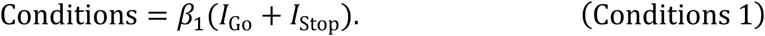

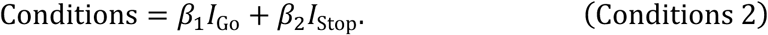

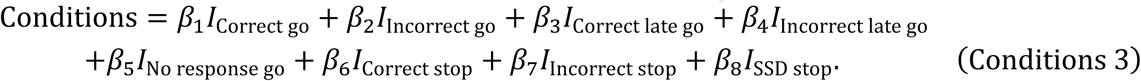

*I*_Condition_ is an indicator function indicating when the subject experiences Condition (for example, *I*_Go_ is 0 except at the moment when a subject is presented with a go trial). For between-subjects analyses, we used Model 1 with Conditions 1 to obtain task activation (β_1_); used Model 1 with Conditions 2 to obtain go activation (β_1_) and stop activation (β_*_); and used Model 1 with Conditions 3 to obtain correct stop versus correct go activation (β_U_ − β_1_), correct stop versus incorrect go activation (β_U_ − β_*_), and incorrect stop versus correct stop activation (β_V_ − β_U_). For correlations between these activations and subject-average behavioral measures, we computed each subject-average measure as the mean of the measure over the trials from both runs during which it assumed values; that is, SSRT was averaged over correct and incorrect stop trials, probability of proactivity and proactive delaying over all trials, and go reaction time over go trials with a recorded response.

To determine within-subjects associations between trial-level brain activity and behavioral measures, for each subject and voxel, we fit the model

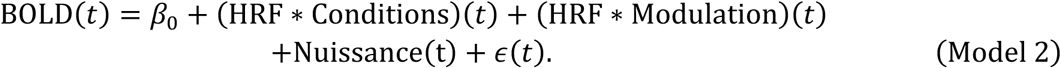

(HRF ∗ Modulation)(*t*) is the value at *t* of the convolution with the hemodynamic response function of the parametric regressor Modulation. To investigate within-subjects associations between brain activity and SSRT on stop trials, probability of proactivity on all trials, proactive delaying on all trials, and observed reaction time on go trials, we set Modulation = β_N_SSRT, Modulation = β_N_P(Proactive), Modulation = β_N_Proactive delaying, and Modulation = β_N_Go RT, respectively, and used Model 2 with Conditions 2. Each of SSRT, *P*(Proactive), Proactive delaying, and Go RT was standardized over the conditions during which it assumed values by subtracting its mean and dividing by its standard deviation; that is, SSRT had mean 0 and standard deviation 1 over correct and incorrect stop trials (and was 0 on all other trials),

*P*(Proactive) and Proactive delaying had mean 0 and standard deviation 1 over all trials, and Go RT had mean 0 and standard deviation 1 over go trials with a recorded response (and was 0 on all other trials).

We fit these regression models for each subject and each of their 2 SST runs. For each model, subject, and voxel, we combined the regression results from the 2 runs with a fixed effects model through FirstLevelModel’s compute_contrast method. For each model and voxel, we estimated the effect of each scanner as the mean of the regression coefficients of the subjects who were scanned by it minus the grand mean of the regression coefficients of all subjects. Then, we adjusted the regression coefficients by subtracting the estimated scanner effects. All analyses of the regression coefficients used these adjusted values. We used a 1-sample Cohen’s *d* (sample mean divided by sample standard deviation) to measure the effect sizes of regression coefficients. Whole-brain correlations and Cohen’s *d* values were visualized on the cortical surface^77^. The surface was obtained using the fetch_surf_fsaverage function in Nilearn’s struct module (mesh=“fsaverage”); data were sampled to the surface using the vol_to_surf function in Nilearn’s surface module; and plotted using the plot_surf_stat_map function in the surf_plotting module.

### Networks and regions of interest

We extracted the whole-brain regression coefficients in 2 networks and 3 sets of regions. We used the Shirer networks for our primary analyses and used the Yeo-17 networks, Shirer regions, cognitive-control regions, and subcortical regions to test the stability of our within-subjects findings. For each subject and each regression coefficient of interest, we obtained the coefficient’s value in each area (network or region) by calculating the mean of the subject’s coefficients over the voxels belonging to the area. We used these area-average regression coefficients of each subject: to compute Cohen’s *d* and Pearson *r* values for the network-level comparison of between- and within-subjects associations (Figure 4) and for the measurement of nonergodicity (Figure 8); and to compute Cohen’s *d* values for the stability (Figure 5) and subgroup (Figure 7) analyses.

The Shirer networks and regions were obtained from ref. ^48^. To obtain the voxel-coordinates of the Yeo-17 networks, we used a mapping between the Brainnetome^78^ and Yeo atlases^79^. We assembled the cognitive-control regions to include 8 areas activated by the SST, 2 core default mode areas, and 1 core salience network and error-processing area. The areas activated by the SST were taken from a meta-analysis of 70 inhibitory control studies^23^ (right anterior insula, right caudate, right inferior frontal gyrus, right middle frontal gyrus, right presupplementary motor area, and right supramarginal gyrus) and a study that segmented high-resolution structural MRI^80^ (left and right subthalamic nucleus). To obtain the default mode areas (posterior cingulate cortex and ventromedial prefrontal cortex), we retrieved a Neurosynth automatic meta-analysis of 777 studies for the term “default mode” on 2024-02-05; extracted clusters from this map using the connected_regions function in Nilearn’s region_extractor module with keyword argument “extract_type” set to “connected_components”; identified by eye the clusters corresponding to the posterior cingulate and ventromedial prefrontal cortex; and for each cluster defined the region to be the 6 mm cube centered on the voxel with the highest meta-analysis Z-score in the cluster. To obtain the salience network and error-processing area (dorsal anterior cingulate cortex), we retrieved a Neurosynth automatic meta-analysis of 464 studies for the term “error” on 2023-09-21 and defined the region to be the 6 mm cube centered on the voxel with the highest meta-analysis Z-score. We obtained subcortical regions from a subcortical probabilistic atlas^81^.

We resampled the atlas’s probabilistic subcortical labels in 1 mm cubed MNI152 2009c nonlinear asymmetric space to the 2 mm cubed MNI152 space of our SPM echo-planar imaging template using resample_to_img from Nilearn’s resampling module, and then thresholded these probabilistic maps at 0.5 to obtain region masks.

### Stability analysis

To assess the stability of the within-subjects results, regression coefficients were resampled at varying sample sizes and the correlation was evaluated against the results in the full sample. Specifically, for each of SSRT, probability of proactivity, and proactive delaying, in each set of networks or regions: 10,000 samples of *n* subjects were drawn with replacement; the Cohen’s *d*’s of each sample’s regression coefficients were calculated and Pearson correlated with the Cohen’s *d*’s of the full sample over the regions or networks; and the mean and 95% and 99% bootstrap confidence intervals were calculated of the correlation (SSRT and probability of proactivity *n* = 25, 40, 70, 120, 200, 335, 560, 945, 1585, 2660, 4469; proactive delaying *n* = 25, 40, 70, 115, 195, 325, 540, 900, 1500, 2505, 4176). The 95% and 99% bootstrap confidence intervals were calculated, respectively, as the intervals covering the 2.5th to 97.5th percentiles and 0.5th to 99.5th percentiles of the 10,000 correlations at each *n*.

We also directly examined the distributions of the Cohen’s *d*’s of the resamples as a function of *n* in regions of interest. Specifically, for each of SSRT, probability of proactivity, and proactive delaying, in each region of interest: 10,000 samples of *n* subjects were drawn with replacement; the Cohen’s *d* of each sample’s regression coefficients was calculated; and the mean and 95% and 99% bootstrap confidence intervals were calculated of the Cohen’s *d* (SSRT and probability of proactivity *n* = 25, 40, 70, 120, 200, 335, 560, 945, 1585, 2660, 4469; proactive delaying *n* = 25, 40, 70, 115, 195, 325, 540, 900, 1500, 2505, 4176). The 95% and 99% bootstrap confidence intervals were calculated as the intervals covering, respectively, the 2.5th to 97.5th percentiles and 0.5th to 99.5th percentiles of the 10,000 Cohen’s *d*’s at each *n*.

### Representational similarity analysis

For each Shirer network, and for each subject, we computed the correlation between the subject’s within-subjects brain maps of SSRT and probability of proactivity, SSRT and proactive delaying, and probability of proactivity and proactive delaying over the voxels in the network. Density estimates used Seaborn’s kdeplot function with each distribution of correlations over subjects normalized to 1 (common_norm=False) and limited to values between -1 and 1 (clip=[-1,1]); all other parameters, including those determining the kernel smoothing bandwidth, were kept at their defaults.

### Measuring nonergodicity of brain networks

For each of SSRT, probability of proactivity, and proactive delaying, and for each of the Shirer networks, we computed bootstrap distributions of a measure of nonergodicity: we drew 10,000 samples of *n* subjects with replacement; for each resample, using the resample’s between- and within-subjects associations, we computed the fraction of subjects whose (within-subjects) brain association with the parameter had the opposite sign of the between-subjects brain association with the parameter, in the network; then, we computed the mean and 95% confidence interval over the resamples of the fraction of opposite signs (SSRT and probability of proactivity *n* = 4469; proactive delaying *n* = 4176). The between-subjects association was the Pearson correlation between correct stop versus correct go activation and the subject-average parameter, which was recomputed for the subjects in each resample. The 95% confidence interval was calculated as the interval covering the 2.5th to 97.5th percentiles of the 10,000 fractions of opposite signs. The goal of using bootstrapping was to account for the strength of the between-subjects results. Next, the nonergodicity of each network was defined as the three-dimensional vector whose *i*th coordinate was the mean (over bootstrap resamples) fraction of subjects with opposite signs in the network for the *i*th cognitive model parameter. Then, the Euclidean distances were computed between the vectors and hierarchical clustering was performed on the distances using the linkage function in scipy’s cluster subpackage, hierarchy module with “method” set to “average”.

### Statistical testing

Permutation tests were used for all statistical testing. The tests used two-sided alternatives and 10,000 resamples and were performed with Scipy’s permutation_test function. FDR correction was performed using the Benjamini-Hochberg procedure with Scipy’s false_discovery_control function. *P*_FDR_ denotes an FDR-corrected *P* value.

For the Shirer networks and an fMRI regression, we tested the null hypothesis for each network that the regression coefficients in the network had a mean of 0 (Figure 4) by computing the means of resamples in which the signs of the coefficients were randomly chosen (permutation_test permutation_type=‘samples’). Then, FDR correction was applied to the *P* values of all networks (e.g., FDR correction was applied to the *P*’s of SSRT’s regression coefficients over the Shirer networks).

For the Shirer networks, an fMRI regression, and a behavioral measure, we tested the null hypothesis for each network that the Pearson correlation between subject-average regression coefficients in the network and subject-average behavioral measures was 0 (Figure 4) by computing the Pearson correlations of resamples in which regression coefficients were randomly paired with behavioral measures (permutation_test permutation_type =‘pairings’). Then, FDR correction was applied to the *P* values of all networks (e.g., FDR correction was applied to the *P*’s of correlations between correct stop versus correct go activation and SSRT over the Shirer networks).

For the Shirer networks and a param γ_1_ or θ_1_, we tested the null hypothesis for each network that the mutually exclusive subgroups of subjects with param < 0 and with param > 0 had different mean regression coefficients (Figure 7) by computing the differences between the means for resamples in which coefficients were randomly assigned to param < 0 and param > (permutation_test permutation_type=‘independent’). Then, FDR correction was applied to the *P* values of all networks (e.g., FDR correction was applied to the *P*’s of mean differences between γ_1_ < 0 and γ_1_ > 0 over the Shirer networks).

For the Shirer networks, we tested the null hypothesis for each network that there was no difference in the network between the median correlation of SSRT and probability of proactivity and the median correlation of SSRT and proactive delaying; the median correlation of SSRT and probability of proactivity and the median correlation of probability of proactivity and proactive delaying; and the median correlation of SSRT and proactive delaying and the median correlation of probability of proactivity and proactive delaying (Figure 6). We computed the differences between the medians of resamples in which correlations were randomly exchanged within subjects (permutation_test permutation_type=‘samples’). Then, FDR correction was applied to the *P* values of all comparisons in all networks. (Since there are 14 Shirer networks and 3 tests per network, FDR correction was applied over 14×3 *P*’s.)

### Software

Data were processed and analyzed using Python (version 3.9.16), Scipy (version 1.11.4), Seaborn (version 0.13.2), Nilearn (version 0.10.1), FSL FLIRT (version 6.0), MATLAB (version R2020b), and JAGS^75^ (version 4.3.0).

