## Supplementary Materials for "Nonergodicity and Simpson’s paradox in neurocognitive dynamics of cognitive control"

### I. Supplementary Results

***Stability of within-subjects brain-behavior associations***

To validate our approach, we investigated the stability and detectability of within-subjects brain-behavior associations in smaller samples, leveraging our large dataset to test the reliability of our findings (**Figure 5**).

To do this, first, we bootstrap resampled the results at varying sample sizes and computed the correlation of these results across brain areas with the results in the full sample using various collections of brain areas. We sampled subjects with replacement, with the number sampled varying between $N = 25$ and the full sample, and then correlated the effect sizes in the resamples with those in the full sample over the areas in each set of networks or regions, with strong correlations demonstrating stability of results across networks and regions. The within-subjects results in samples as small as 25 subjects correlated strongly with the results in the full sample.

Second, we directly examined the distributions of the effect sizes in the resampled data for regions of interest to gain another perspective on the reliability of the within-subjects findings. These finer-grained consistency checks showed that key results were observed frequently in small and moderate samples. For example, proactivity, as measured by either proactive parameter, was associated with right anterior insula suppression in greater than 95% of $N = 25$ samples, while proactivity's association with activation in the left retrosplenial cortex required $N \sim 100$ to emerge in greater than 95% of samples. However, some results showed considerable variability in the direction of effects for samples of $N > 100$, for example SSRT in the right anterior insula and proactive delaying in the right retrosplenial cortex.

These analyses demonstrate that within-subjects associations between brain activity and our model parameters (SSRT, probability of proactivity, and proactive delaying) were stable and detectable even with modest sample sizes. While some effects required larger samples to emerge consistently, many key findings were prevalent even in small samples. This stability supports the validity of our within-subjects approach and suggests that meaningful insights into neurocognitive mechanisms can be gained from studies with more modest sample sizes.

***Robustness of nonergodicity to analytical choices***

To test the robustness of our nonergodicity findings, we conducted several control analyses. These analyses aimed to evaluate whether nonergodicity in brain-behavior relations was specific to the brain and behavioral measures used in the main analyses.

<u>Alternative between-subjects analyses</u>

Our primary analysis compared within-subjects and between-subjects brain-behavior associations using a canonical approach for the SST[1-6]: correlating subject-average behavioral measures with subject-average differences in brain activation between correct stop and correct go trials. However, other between-subjects analyses can also be reasonably compared against the within-subjects analyses. We tested three alternative between-subjects approaches, each correlating the subject-average parameters (SSRT, probability of proactivity, and proactive delaying) with different subject-average brain measures from the task. First, to align more closely with our within-subjects analyses, we (i) correlated subject-average SSRT with subject-average brain activation from all stop trials (correct and incorrect combined), and (ii) correlated subject-average probability of proactivity and proactive delaying with subject-average brain activation from all trials (go and stop combined) (**Supplementary Figure S2a**). Second, we used two other standard subject-average brain measures[4]: (i) difference in activation between incorrect stop and correct go trials (incorrect stop versus correct go activation) (**Supplementary Figure S2b**) and (ii) difference in activation between incorrect and correct stop trials (incorrect stop versus correct stop activation) (**Supplementary Figure S2c**). Comparing these between-subjects results to the within-subjects results (**Supplementary Figure S2d**) demonstrates that nonergodicity for the cognitive model parameters persisted across different analytical choices.

Nonergodicity of brain associations with an observed behavioral measure

To determine whether nonergodicity was specific to latent model-derived parameters or a general feature of brain-behavior associations in this task, we repeated our analyses using go reaction time, a directly observed behavioral measure. Between-subjects, we correlated subject-average go reaction time with correct stop versus correct go activation (**Supplementary Figure S3a**), incorrect stop versus correct go activation (**Supplementary Figure S3b**), incorrect stop versus correct stop activation (**Supplementary Figure S3c**), and go activation (subject-average activation from all go trials) (**Supplementary Figure S3d**). Within-subjects, we examined the association of brain activity with reaction time on go trials (**Supplementary Figure S3e**). Comparison of these between- and within-subjects results indicated widespread nonergodicity of this observed behavioral measure's brain associations.

In all, we examined 4 different strategies of performing between- and within-subjects analysis of how 3 latent cognitive parameters and 1 observed behavioral measure related to brain activity. In each of these 16 comparisons, within- and between-subjects inferences about brain-behavior associations diverged. This consistent finding across multiple analytical approaches and measures provides robust evidence that the neurocognitive dynamics of inhibitory control are fundamentally nonergodic.

### II. Supplementary Figures

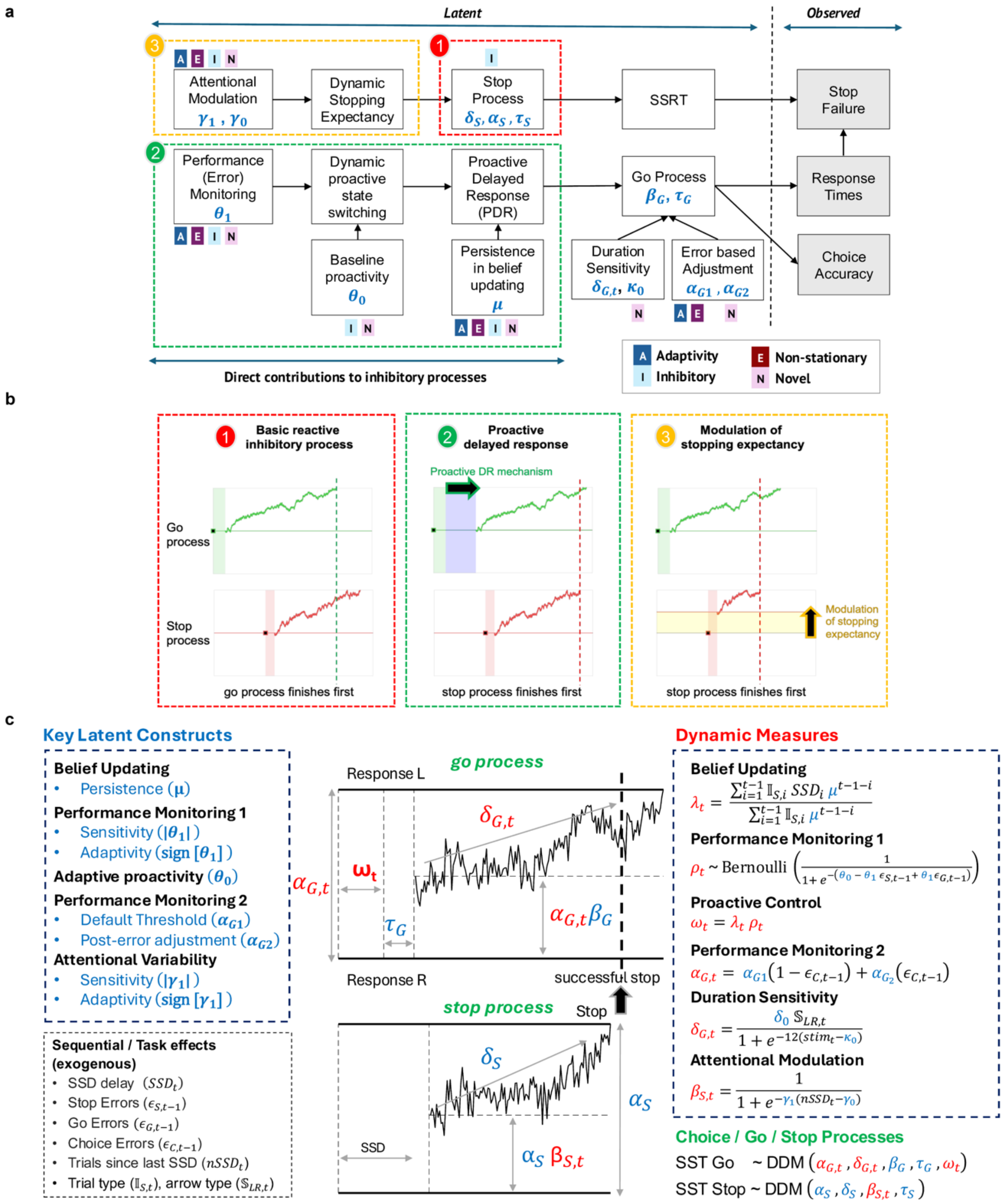


**Supplementary Figure S1. The PRAD cognitive model. a,** The PRAD model infers latent variables for each subject from their observed go and stop failure rates, response times, and choice accuracy. The latent variables relate to three mechanisms of dynamic inhibitory control: the basic reactive inhibitory process (red 1), proactive delaying of responses (green 2), and modulation of stopping expectancy (yellow 3). **b,** Visualization of the three mechanisms of dynamic inhibitory control. **c,** Mathematical details of how the model parameterizes the go and stop processes.

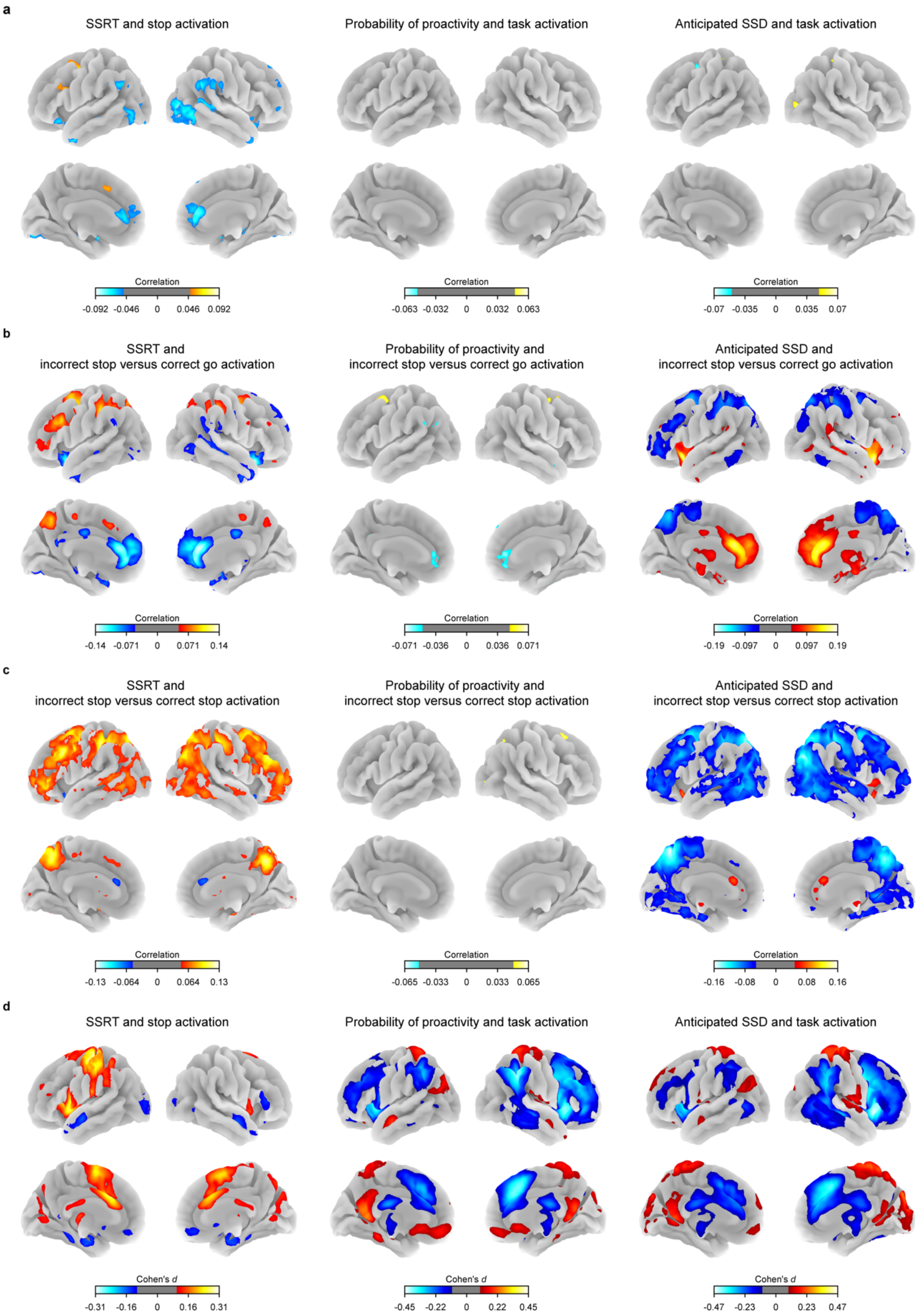


**Supplementary Figure S2. Nonergodicity is robust to type of between-subjects analysis. a-c,** Between-subjects correlation maps of associations between SSRT and stop activation, probability of proactivity and task activation, and proactive delaying and task activation **(a)**; and between-subjects correlation maps of associations between the cognitive model parameters and incorrect stop versus correct go activation **(b)**, and incorrect stop versus correct stop activation **(c)**. In each voxel, subject-average brain activation was correlated with subject-average SSRT, probability of proactivity, and proactive delaying. The correlation maps were thresholded at $\geq 0.05$. **d,** Within-subjects Cohen's $d$ maps of associations between cognitive model parameters and brain activity. For each subject and in each voxel, brain activity was regressed on: SSRT on stop trials, probability of proactivity on all trials, and proactive delaying on all trials. The resulting Cohen's $d$ maps were thresholded at $\geq 0.1$.

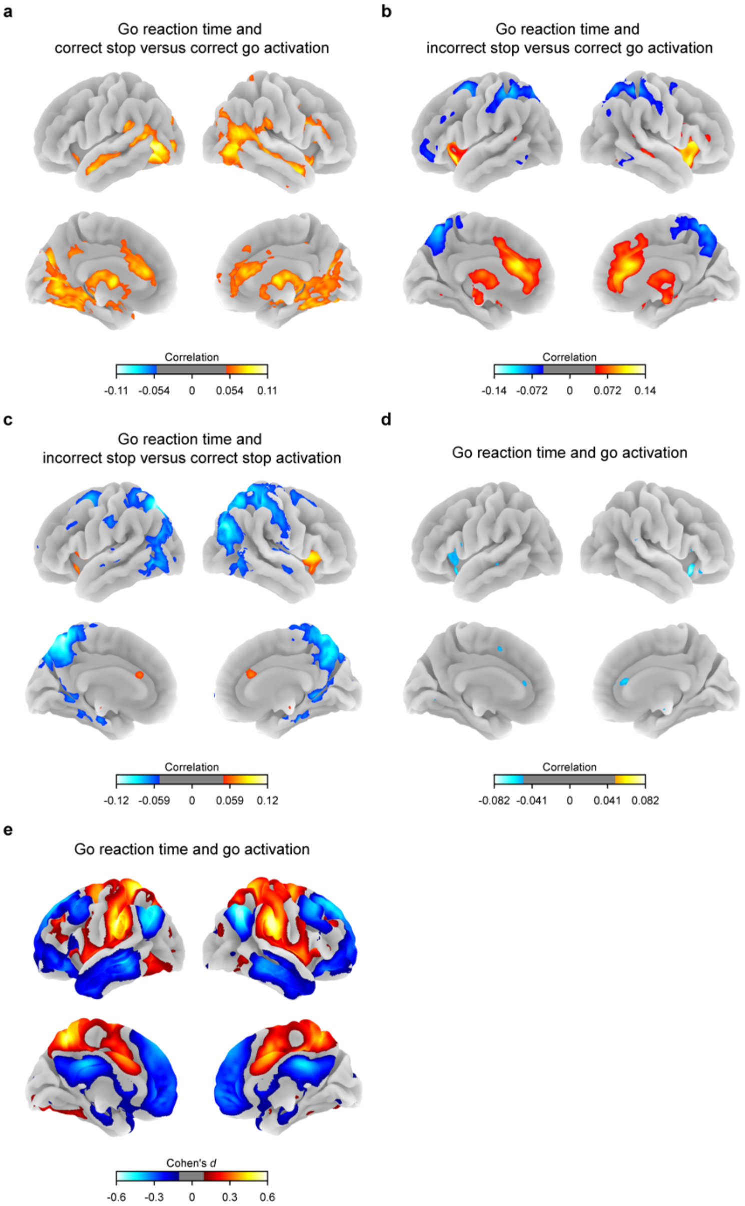


**Supplementary Figure S3. Nonergodicity is exhibited by directly observed behavioral measure. a-d,** Between-subjects correlations maps of associations between go reaction time and correct stop versus correct go activation **(a)**, incorrect stop versus correct go activation **(b)**, incorrect stop versus correct stop activation **(c)**, and go activation **(d)**. In each voxel, subject-average go reaction time was correlated with subject-average brain activation. The correlation maps were thresholded at $\geq 0.05$. **e,** Within-subjects Cohen's $d$ map of associations between go reaction time and brain activity. For each subject and in each voxel, brain activity was regressed on reaction time on go trials; the Cohen's $d$'s of the regression coefficients were then calculated. The resulting Cohen's $d$ map was thresholded at $\geq 0.1$.
